# Decoding Heterogenous Single-cell Perturbation Responses

**DOI:** 10.1101/2023.10.30.564796

**Authors:** Bicna Song, Dingyu Liu, Weiwei Dai, Natalie McMyn, Qingyang Wang, Dapeng Yang, Adam Krejci, Anatoly Vasilyev, Nicole Untermoser, Anke Loregger, Dongyuan Song, Breanna Williams, Bess Rosen, Xiaolong Cheng, Lumen Chao, Hanuman T. Kale, Hao Zhang, Yarui Diao, Tilmann Bürckstümmer, Jenet M. Siliciano, Jingyi Jessica Li, Robert Siliciano, Danwei Huangfu, Wei Li

**Affiliations:** Center for Genetic Medicine Research, Children’s National Hospital, Washington DC, USA; Department of Genomics and Precision Medicine, George Washington University, Washington DC, USA; Developmental Biology Program, Sloan Kettering Institute, New York City, NY, USA; Louis V. Gerstner Jr. Graduate School of Biomedical Sciences, Memorial Sloan Kettering Cancer Center, New York City, NY, USA; Department of Medicine, Johns Hopkins University School of Medicine, Baltimore, MD, USA; Howard Hughes Medical Institute, Johns Hopkins University School of Medicine, Baltimore, MD, USA; Department of Statistics and Data Science, University of California, Los Angeles, CA, USA; Myllia Biotechnology GmbH. Am Kanal 27, 1110 Vienna Austria; Bioinformatics Interdepartmental Ph.D. Program, University of California, Los Angeles, CA, USA; Weill Cornell Graduate School of Medical Sciences, Weill Cornell Medicine, New York, NY, USA; Department of Cell Biology, Duke University Medical Center, Durham, NC, USA; Department of Human Genetics, University of California, Los Angeles, CA, USA; Department of Biostatistics, University of California, Los Angeles, CA, USA; Department of Computational Medicine, University of California, Los Angeles, CA, USA

**Keywords:** Perturb-seq, CRISPR-based genetic perturbations, single-cell RNA-seq, computational model

## Abstract

Understanding diverse responses of individual cells to the same perturbation is central to many biological and biomedical problems. Current methods, however, do not precisely quantify the strength of perturbation responses and, more importantly, reveal new biological insights from heterogeneity in responses. Here we introduce the perturbation-response score (PS), based on constrained quadratic optimization, to quantify diverse perturbation responses at a single-cell level. Applied to single-cell transcriptomes of large-scale genetic perturbation datasets (e.g., Perturb-seq), PS outperforms existing methods for quantifying partial gene perturbation responses. In addition, PS presents two major advances. First, PS enables large-scale, single-cell-resolution dosage analysis of perturbation, without the need to titrate perturbation strength. By analyzing the dose-response patterns of over 2,000 essential genes in Perturb-seq, we identify two distinct patterns, depending on whether a moderate reduction in their expression induces strong downstream expression alterations. Second, PS identifies intrinsic and extrinsic biological determinants of perturbation responses. We demonstrate the application of PS in contexts such as T cell stimulation, latent HIV-1 expression, and pancreatic cell differentiation. Notably, PS unveiled a previously unrecognized, cell-type-specific role of coiled-coil domain containing 6 (CCDC6) in guiding liver and pancreatic lineage decisions, where CCDC6 knockouts drive the endoderm cell differentiation towards liver lineage, rather than pancreatic lineage. The PS approach provides an innovative method for dose-to-function analysis and will enable new biological discoveries from single-cell perturbation datasets.

**One sentence summary:** We present a method to quantify diverse perturbation responses and discover novel biological insights in single-cell perturbation datasets.

## Introduction

Perturbation is essential for understanding the functions of the mammalian genome that encodes proteincoding genes and non-coding elements (e.g., enhancers). Single-cell profiling of cells undergoing genetic, chemical, environmental or mechanical perturbations is commonly used to examine perturbation responses at the single-cell level. Recently, high-throughput approaches of perturbation have been developed using single-cell RNA-seq (scRNA-seq) readout, including multiplexing of perturbations and single-cell CRISPR screen (e.g., Perturb-seq, CROP-seq)^1–7^. This concept has been extended to study changes in single-cell chromatin accessibility^8,9^, spatial transcriptomics^10^ upon perturbations or perturbation combinations^11–13^, and other phenomena.

Understanding how perturbations lead to different responses within cells is critical to understanding the fundamental biology behind perturbation. Technical factors, including single-cell assays used to profile the response, and the on-target/off-target effects of perturbations, are known drivers that lead to differences of single-cell profiles in the data^14–16^. In Perturb-seq experiments that use CRISPR/Cas9 for knockouts, both in-frame deletions^16^ and chromosomal losses^17^ contribute to different expression profiles and clustering patterns of single cells.

Perhaps more interestingly, the heterogeneity perturbation responses may be driven by underlying biological factors (**Fig. 1a**). These factors may be either cell-intrinsic (e.g., the activities of other coding- and non-coding genomic elements) or cell-extrinsic (e.g., cell states or types, environment factors), all of which define the context of perturbation response. For example, combined expressions of transcription factors (TFs) are critical for many cellular state conversions. Therefore, to properly decode the functions of these TFs via perturbation, one must consider the effect of the cell state and the activities of other companion TFs. For this reason, defining the heterogeneity of perturbation response and identifying factors that contribute to these outcomes is important for understanding how cells respond to perturbation.

**Figure 1.**
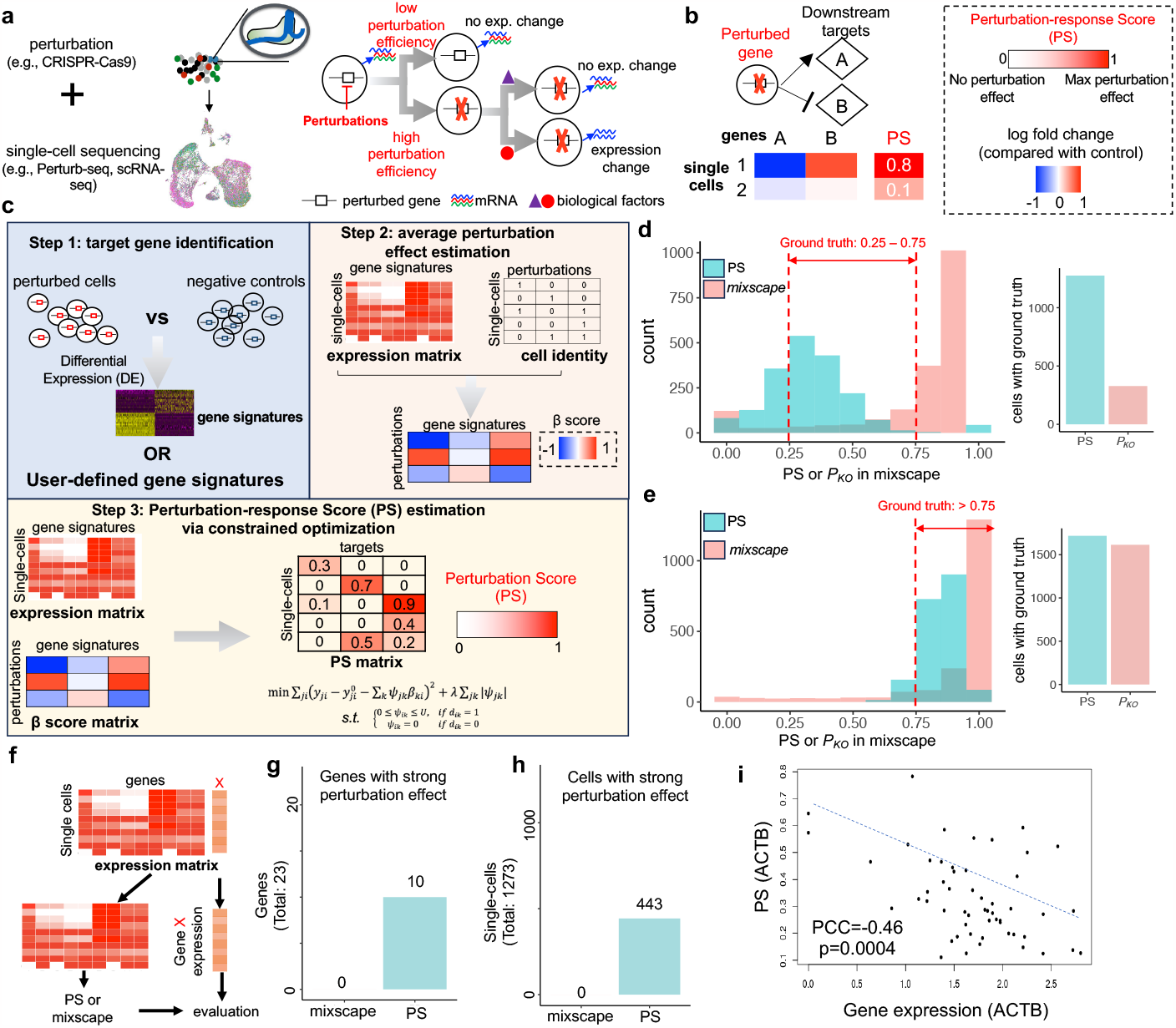
The Perturbation-response Score (PS) framework and benchmark. **a**, Overview of different technical and biological factors that contribute to heterogenous perturbation outcomes from single-cell perturbation datasets. **b**, Using downstream gene expressions to infer the value of PSs. **C**, Overview of the scMAGeCK-PS that estimates PS value. **d-e**, Benchmark results of both PS and mixscape using simulated datasets, where 50% (**d**) and 100% € gene perturbation effects are simulated using scDesign3. Here, the expressions of 200 differentially expressed genes (DEGs) from bulk RNA-seq (Nelf knockout vs. wild-type) are simulated, and ground truth efficiency value is indicated in red color. **f**, Benchmark pipeline using real CRISPRi-based Perturb-seq datasets, where the perturbation efficiency can be evaluated directly via gene expression. **g-h**, Benchmark results of mixscape and scMAGeCK-PS using a published Perturb-seq dataset, by counting the numbers of cells or genes with strong perturbation effects. A gene is considered to have strong perturbation effect, if a strong negative correlation (Pearson correlation coefficient < -0.1) is observed between PS and the expression of that gene across all perturbed cells. A cell is considered to be strongly perturbed, if its predicted efficiency score (by scMAGeCK-PS or mixscape) within one cell is greater than 0.5. The Perturb-seq experiment is performed with low MOI condition, where most cells have only 1 expressed guide. **i**, An representative estimation results of scMAGeCK-PS and their correlations of ACTB expression.

Unfortunately, computational frameworks are currently lacking to decode the diverse outcomes of perturbations. For technical factors, mixscape is the only method to detect and mitigate confounding variations (e.g., incomplete knockouts from CRISPR/Cas9)^16^. However, its performance has not been rigorously benchmarked, especially when partial gene functions are perturbed using techniques like CRISPR interference (CRISPRi). More importantly, no methods have been developed to reveal new biological insights from the heterogenous perturbation outcomes, including studying how partial gene perturbations affect a phenotype of interest (i.e., “dosage” analysis), and discovering biological determinants that govern differential perturbation responses.

Here we present a computational framework, the perturbation-response score (PS), to quantify heterogenous perturbation outcomes in single-cell transcriptomics datasets. The PS, estimated from constrained quadratic optimization, quantifies the strength of the perturbation outcome for a single cell. We performed comprehensive benchmark studies that demonstrated the outstanding performance of PS over existing methods, including simulated datasets, genome-scale Perturb-seq, and published Perturb-seq datasets that cover various CRISPR-based technologies. More importantly, PS analysis presents two major conceptual advances for analyzing single-cell perturbation data: it enables analysis of the dose of perturbation, and identification of novel biological determinants that govern the heterogeneity of perturbation responses. First, we analyzed essential gene Perturb-seq and found two patterns of dose response, based on whether moderate perturbation leads to strong expression changes of downstream genes. Second, we identified intrinsic and extrinsic biological factors governing critical gene functions in latent HIV-1 expression and pancreatic/liver development. Based on PS analysis results, we identified and confirmed a novel function of CCDC6, wherein perturbation drives duodenum cell differentiation towards liver commitment. Collectively, PS analysis provides a powerful tool to decode heterogenous perturbation outcomes from single-cell assays.

## Results

### Using PS to detect heterogenous perturbation outcomes within and across datasets

Perturbing the same gene (or non-coding elements) may result in different phenotypic changes or transcriptional outcomes (**Fig. 1a**), depending on technical factors (e.g., perturbation efficiency) and biological factors (e.g., cell type, cell state, activities of cofactors). Unfortunately, existing methods can detect only technical factors^16^, while biological factors remain unexplored. To bridge this gap, we built a computational framework to quantify perturbation outcomes in single-cell datasets using scRNA-seq as readout. Corresponding assays include single-cell CRISPR screens (e.g., Perturb-seq), or simply multiplex scRNA-seq profiling of various perturbations (e.g., sci-Plex; **Fig. 1b, c**). We define the perturbation-response score (PS) to quantify the strength of perturbation, where PS=0 indicates no perturbation effect (e.g., effects corresponding to unperturbed, wild-type gene functions) and PS=1 indicates the maximum perturbation effect observed within a dataset; for example, effects that correspond to homozygous knockouts on both alleles of a gene. We utilize the expressions of multiple downstream targets of a perturbed gene to infer the (unknown) values of PS (**Fig. 1b**). For example, if one cell has dramatic expression changes on the known downstream target genes, then its value of PS should be higher than cells with weak expression changes of these genes.

We built a computational model, based on a constrained quadratic optimization, to automatically identify the downstream targets of perturbed genes and calculate PS (**Fig. 1c**). This model, named “scMAGeCK-PS”, is based on our previously published scMAGeCK algorithm^15^ and consists of three steps. First, scMAGeCK-PS identifies differentially expressed genes (DEGs) upon perturbation (e.g., perturbing the function of gene *X*), by comparing the transcriptome profiles between perturbed cells and unperturbed cells. These DEGs are served as “signature” target genes of *X*. Second, scMAGeCK-PS used a previously developed scMAGeCK model to estimate the average effect of perturbation on these target genes, which can be estimated from the first step. Third, scMAGeCK-PS uses a constraint optimization procedure to find the value of PS that minimizes the sum of mean squared errors between predicted and measured expression changes of all downstream targets (see Methods). The constraints are established such that any PS is non-negative for cells with *X* perturbed, and is exactly zero in cells without perturbation. Such constraints can be established based on the prior information of perturbations; for example, the expression matrix of single-guide RNAs (sgRNAs).

### PS outperforms mixscape in quantifying partial perturbations

mixscape^16^ is currently the only method to detect and remove technical factors that affect perturbation outcomes, especially incomplete gene knockouts that are generated from CRISPR/Cas9. However, the performance of mixscape on partial gene perturbations has not been fully evaluated. Here we compare PS with mixscape using multiple benchmark datasets. We first used synthetic datasets to evaluate the performances of different methods, because finding a real scRNA-seq dataset that contains ground truth (*i*.*e*., accurate measurements of loss-of-function upon perturbation) is challenging. For synthetic data generation, we used scDesign3^18^ to simulate the single-cell transcriptomic responses upon perturbing the 50% and 100% functions of Nelfb, based on a real scRNA-seq dataset that deletes Nelfb in mouse T cells^19^ (**Supplementary Fig. S1a;** see Methods). We specified different numbers of DEGs (from 10 to 500) and simulated their expression changes upon 50% or 100% perturbations of Nelfb functions. In all the cases, PS correctly estimated partial perturbation, where the median PSs range from 0.32-0.34 for 50% perturbation, and greater than 0.8 for 100% perturbation, respectively (**Fig. 1d-e; Supplementary Fig. S1b-e**). In contrast, mixscape uniformly assigned the posterior probability of perturbation to 1 in all cases, an indication that mixscape is not suited to analyze the outcome of partial gene perturbations (**Fig. 1d-e; Supplementary Fig. S1b-e)**, possibly due to the bimodal statistic model it uses, which only considers 100% knockout effects^16^.

We next evaluated different methods using real single-cell perturbation datasets. We chose CRISPRi-based Perturb-seq datasets (**Fig. 1f-i**) because the CRISPRi system directly modulates the expression levels of perturbed genes, and the perturbation efficiency can be accessed using single-cell transcriptomic data. We use two published K562 CROP-seq datasets:^20^ in the first, only 1 gRNA is expressed within each cell (*i*.*e*., low multiplicity of infection or MOI), and in the second multiple gRNAs are expressed (*i*.*e*., high MOI). We examine cells where the transcription starting sites (TSS) of highly expressed protein-coding genes are targeted (23 and 342 genes, respectively; **Fig. 1g-i, Supplementary Fig. S1f-g**). If the TSS of gene *X* is perturbed, we first removed the expression of *X* from expression matrix, and used the rest of gene expressions to measure the perturbation efficiency of *X*. The scores of different methods were then compared with the expression of *X*, producing a direct measurement of perturbation efficiency (**Fig. 1f**). In over 40% of these genes (10 out of 23 for low MOI, 139 out of 342 for high MOI), PS has a strong negative correlation with the expression of *X* (**Fig. 1g**), defined as Pearson correlation coefficient < -0.1 and p value <0.01. In contrast, mixscape scores correlate with X expression in none of the genes (for low MOI dataset; **Fig. 1g)**, or in less than 5% of all the genes (for high MOI dataset; **Supplementary Fig. S1f-g**). PS detects a much greater number of cells that have a strong perturbation effect (PS or mixscape score >0.5; **Fig. 1h**), whose scores are strongly negatively correlated with gene expression (**Fig. 1i**). We also tested both methods in another CRISPRi-based Perturb-seq dataset, where sgRNAs with mismatches reduce efficiency, leading to partial perturbation effects^21^ (**Supplementary Fig. S1h-i**). PS has a high sensitivity and a good balance between sensitivity and specificity, evidenced by the higher Pearson correlation coefficients (**Supplementary Fig. S1h**) and areas under the receiver-operating characteristic (ROC) curve (AUC) values (**Supplementary Fig. S1i**).

To further benchmark methods in terms of a phenotype of interest, we designed and performed a genome-scale CRISPRi Perturb-seq on both unstimulated and stimulated Jurkat, a T lymphocyte cell model (**Fig. 2a**), and evaluated the performances of different methods in identifying known regulators of T cell activation. We designed Perturb-seq library that contains sgRNAs targeting the TSS of 18,595 genes (4-6 guides per gene) and used a TAP-seq-based^22^ multiplex primer panel to detect the expressions of 374 genes with high sensitivity (see **Supplementary Table S1** and Methods). We obtained high-quality scRNA-seq data on over 586,000 single cells after quality control, and the UMAP clustering of Perturb-seq datasets clearly demonstrated the differences between stimulated and non-stimulated cells (**Fig. 2b**). Next, we ran PS or mixscape to calculate the scores of all perturbations at a single-cell level; and for each perturbed gene, we calculated its overall perturbation-response score, by adding the scores of all cells that express a corresponding sgRNA targeting that gene. Because our system focuses on T cell stimulation, perturbing a gene that reaches a highest (and lowest) cumulative score should have the strongest (and no) effect on T cell stimulation, respectively. For an independent evaluation, we extracted 385 (and 1297) positive (and negative) hits whose perturbation impairs (or does not impair) the stimulation of T cells from a published genome-scale CRISPR screen^23^. Both Perturb-seq and pooled CRISPR screen identified many known positive regulators of T cell activation, such as components of the T cell receptor complex (e.g., CD3D) and proximal signaling components (e.g., LCK; **Fig. 2c**). For many positive genes, cells with higher values of PS or mixscape score are skewed towards non-stimulating state, consistent with their negative selections in pooled CRISPR screens using T cell stimulation as readout (**Fig. 2c; Supplementary Fig. S2**). However, when comparing the ROC score, PS reaches a higher AUC score than mixscape (**Fig. 2d**), indicating its better performance in accurately separating positive from negative hits.

**Figure 2.**
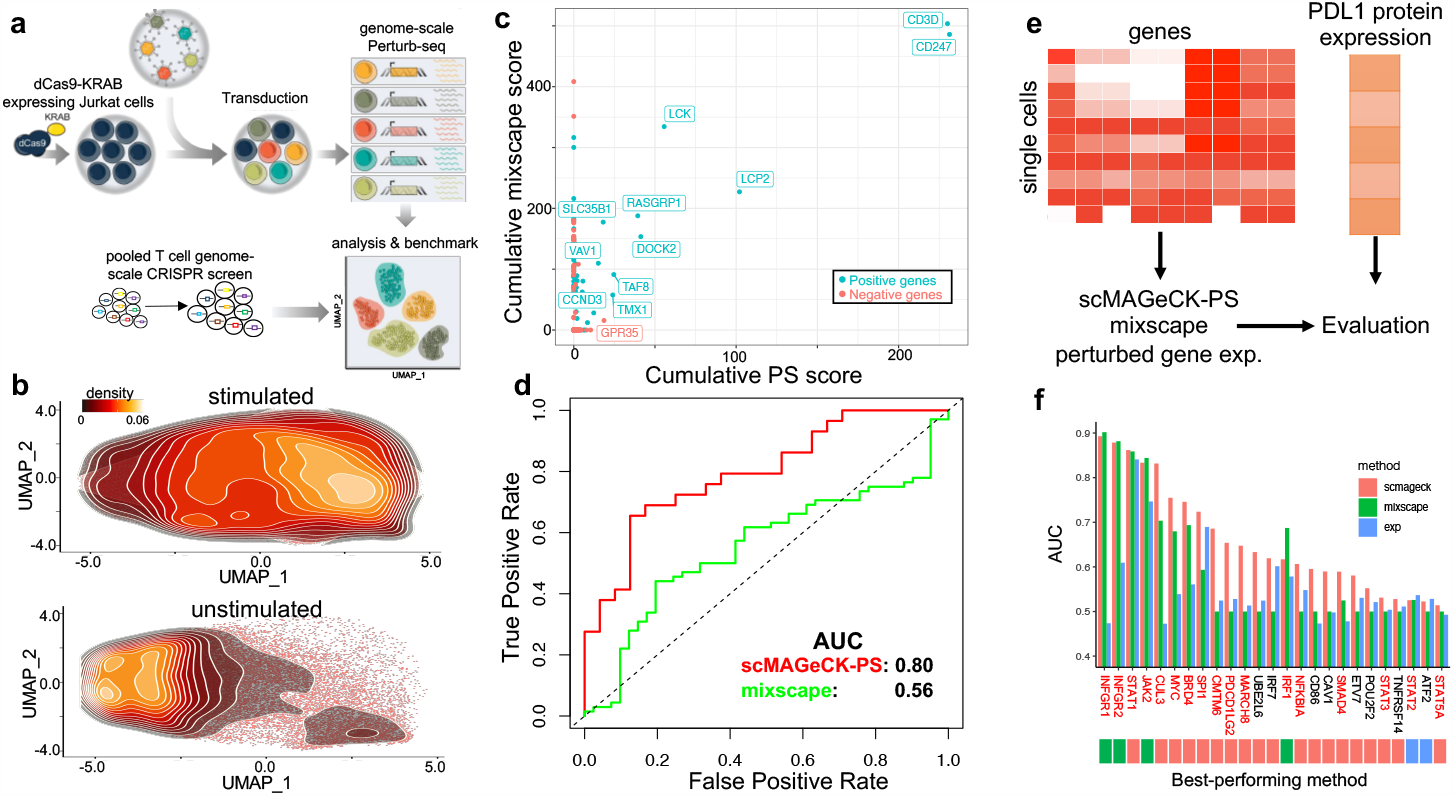
Additional benchmark results using genome-scale Perturb-seq and ECCITE-seq. **a**, Benchmark procedure using a genome-scale Perturb-seq and a published, pooled T cell CRISPR screen. **b**, The distribution of unstimulated and stimulated Jurkat cells along the UMAP plot. **c**, The correlation of predicted scores by scMAGeCK-PS and mixscape. **d**, the Receiver-Operating Characteristic (ROC) curve of both methods in separating positive and negative hits. **e-f**, Benchmark using a published ECCITE-seq where PDL1 protein expression is used as gold standard (**e**), and the performance of different methods in terms of predicting PLD1 protein expression (**f**).

Finally, we tested different methods on a published ECCITE-seq, which simultaneously measures single-cell transcriptomes, surface proteins, and perturbations^16^. PDL1 protein expression was used as an independent metric of evaluation (**Fig. 2e**), because PDL1 is a well-studied gene whose protein expression is well understood. Among 25 perturbed genes in the ECCITE-seq perturbation library, 17 are known to regulate PDL1 expression (**Fig. 2f**). We compared PS with mixscape in terms of predicting changes in PDL1 expression (**Fig. 2f; Supplementary Fig. S3)**. In addition, the expression of the perturbed gene is included in the comparison as a naïve method. In 19 out of 25 genes (76%), PS outperformed mixscape and perturbed gene expression in predicting PDL1 expression (**Fig. 2e**), including 12 out of 17 (71%) known PDL1 regulators. Notably, for genes whose perturbations led to strong transcriptomic changes (*e*.*g*., IFNGR1, IFNGR2, JAK2, STAT1), both PS and mixscape work well, reaching AUC > 0.8 (**Fig. 2f**). For other genes whose perturbation only leads to moderate or weak expression changes, as described previously^16^, PS outperforms mixscape, including those that are confirmed to be PLD1 regulators (*i*.*e*., genes marked in red in **Fig. 2f**).

### Analyzing dose-dependent effects of perturbation

Traditionally, dosage analysis requires a careful, time-consuming adjustment of perturbation strength, including changing drug concentrations or designing sgRNA sequences to achieve various editing efficiencies^21,24^. Since the quantifying partial gene perturbation by PS is highly accurate (**Fig. 1-2**), we can use PS to perform dose-response analysis of perturbation, without the need to titrate the strength of perturbation. By examining ECCITE-seq data in which PDL1 expression was measured directly (**Fig. 2e**), we found correlations between PDL1 expression and the PS of known PDL1 regulators (**Fig. 3a**). The PSs of positive PDL1 regulators (e.g., IFNGR1/2, STAT1; **Fig. 3b**) are negatively correlated with PDL1 expression, while the scores of negative regulators (e.g., CUL3, BRD4) are negatively correlated (**Supplementary Fig. S3; Fig. 3c**). One example is CUL3, which is known to destabilize and degrade PDL1 protein expression^25^. Consequently, higher CUL3 PSs, indicating higher CUL3 functional perturbation, correspond to higher PDL1 protein expressions (**Fig. 3a**). Compared with mixscape, PS more accurately predicts the quantitative changes in PDL1 expression, evidenced by stronger Pearson correlations between the two (**Supplementary Fig. S3**).

**Figure 3.**
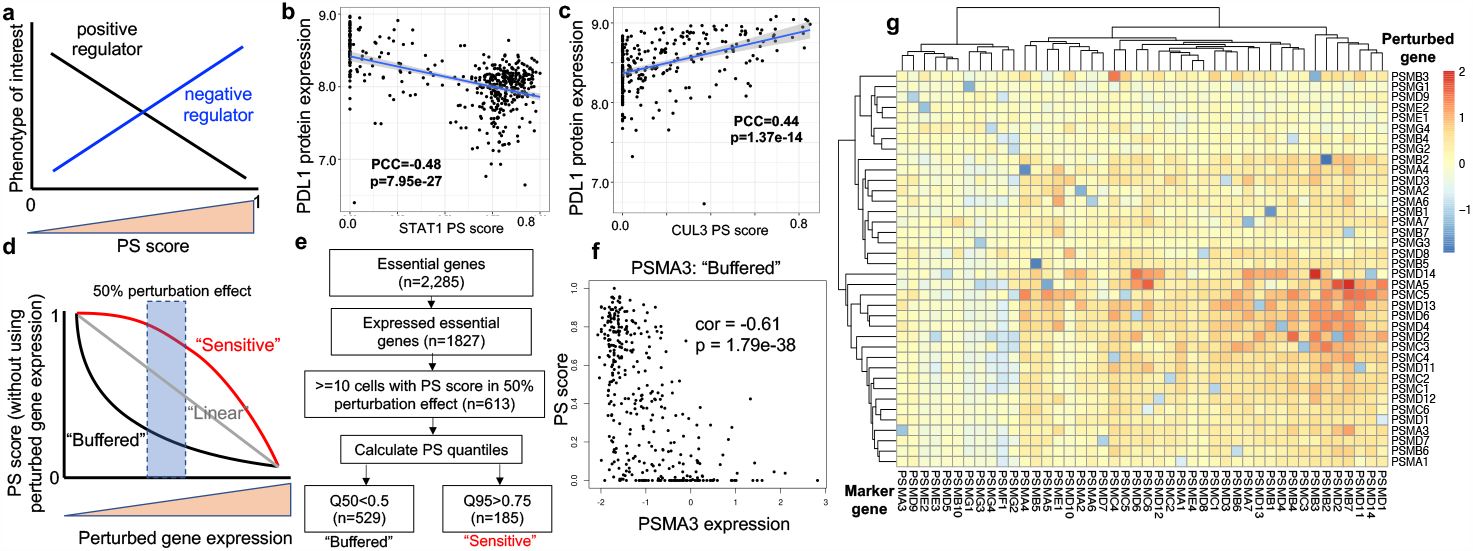
Dose-dependent responses of perturbations. **a**, The correlation between a gene’s PS and a phenotype of interest indicates positive (or negative) regulations. **b-c**, The correlation between PDL1 protein expression and the PS of CUL3 (**b**) and STAT1 (**c**). CUL3 is a known negative regulator of PDL1, while STAT1 is a known positive regulator. **d**, The classification of buffered or sensitive genes, based on perturbed gene expression and PS. **e**, The classification of buffered or sensitive genes from published Perturb-seq datasets focusing essential genes in K562^26^. **f**, The perturbation-expression plot of PSMA3, a buffered gene. **g**, The log fold changes of mark gene expressions (columns) upon perturbing proteasome genes (rows) from the essential gene Perturb-seq dataset.

We further investigated the relationships between perturbation efficiency and the strength of perturbation responses, which is measured by PS (**Fig. 3d**). In particular, we are interested in genes that show one of two different patterns of PSs upon perturbation: “buffered” distribution, where genes have high PSs only when stronger perturbation efficiency is achieved; and “sensitive” distribution, where the PSs are high, even with moderate or weak perturbation efficiency. Both “buffered” or “sensitive” terms have been coined previously to describe the effects of transcription factor dosages to chromatin accessibility^26^. CRISPRi-based Perturb-seq datasets are used, as the efficiencies of CRISPR inhibition can be directly evaluated by examining perturbed gene expressions (**Fig. 2e**).

We calculated PS for every gene in a published essential-wide Perturb-seq^27^, which uses CRISPRi to inhibit the expressions of 2,285 common essential genes. We classified genes based on their PS quantiles that correspond to around 50% perturbation efficiency (**Fig. 3e**): a gene is classified as “buffered” if its median PS is smaller than 0.1; or “sensitive”, if its 95% quantile is greater than 0.75. Among over 2,000 essential genes, we classified 613 genes as either buffered or sensitive. The majority are buffered (529 out of 613), indicating high robustness to perturbation, possibly due to their essential roles in cellular functions that require compensations on expression reductions. Many buffered genes belong to essential protein complexes, including proteosomes (e.g., PSMA3; **Fig. 3f**) and ribosomal subunits (e.g., RPL4; **Supplementary Fig. S4a**). 30% of the genes (185 out of 613) belong to “sensitive” category, showing strong transcriptome responses even with moderate or weak efficiencies upon perturbing gene expression (**Supplementary Fig. S4b-c**). Many of the sensitive genes are also displaying buffering effect, a demonstration of complex, heterogenous responses of cells undergoing the same perturbation of essential genes. Notably, 50% reduction of HSPA5 and GATA1 expression achieved near-maximal transcriptional response (and the associated growth defect), as in previous studies^22^.

We further examined possible mechanisms by which buffered genes resist perturbation, especially those that belong to the same functional protein complex. Interestingly, perturbing one member of the protein complex usually leads to the expression up-regulation of other members of the complex, indicating a possible mechanism for compensation. For example, perturbing proteosome subunits led to a strong expression reduction of the perturbed gene (*e*.*g*., PSMA5; blue squares in **Fig. 3g**) and concurrent up-regulation of other members of the proteosomes (*e*.*g*., PSMB7, PSMD2). Perturbing many other protein complexes, including ribosomal subunit, mediator, and RNA polymerases, also leads to similar up-regulation of some members of the same functional unit (**Supplementary Fig. S5a-c**), indicating that compensation occurs by up-regulation of other submits of the same molecular machine. To confirm our findings on a different cellular system, we examined the effects of perturbing proteosomes in our genome-scale Perturb-seq dataset (**Fig. 2a**). The TAP-seq approach used in this dataset provides a sensitive and accurate measurement of gene expression changes upon perturbation^27^. Indeed, perturbing members of the proteosome subunits leads to the up-regulation of other proteosomes (**Supplementary Fig. S5d**), consistent with the known transcriptional feedback loop that is observed between proteosome genes^28^. Overall, the widespread existence of such compensatory effect may explain the perturbation-expression phenotype of buffered genes, where a strong perturbation efficiency is needed to achieve strong expression changes.

### PS reveals intrinsic and extrinsic biological factors that regulate gene functions in latent HIV expression

We next perform Perturb-seq experiment and use PS to investigate the functions of key genes regulating latent HIV-1 expression. We used a Jurkat HIV cell model that we previously established for pooled CRISPR screening^29^, where cells stably express Cas9 and are latently infected with HIV-GFP viral vector. We designed a Perturb-seq library that targets 10 protein-coding genes (**Supplementary Table S2**), which are either (1) known factors in HIV-1 virus expression and T cell activation (e.g., BIRC2), or (2) top hits from genome-scale CRISPR screens that we previously performed (e.g., BRD4)^29^. We performed Perturb-seq experiments in three different conditions, including stimulated Jurkat (by PMA/I) followed by GFP expression sorting (GFP+ or GFP-), and unstimulated cells (**Fig. 4a**). The single-cell transcriptomes were profiled via the 10X Genomics Chromium platform, and expressed guide RNAs can be captured directly. After quality controls, we received 7,063-8,811 single cells per sample, where the mean reads per cell (and median genes expressed per cell) in each sample is at least 69,888 (and 4,744), respectively (**Supplementary Fig. S6a**). Guide RNAs were detected in over 96% of the cells, and over 85% of these cells could be assigned a unique guide RNA (**Supplementary Table S3**). The transcriptome profiles of cells are primarily clustered by cell states (stimulated vs. unstimulated), indicating that the primary sources of expression variation are coming from cell states (**Fig. 4b**).

**Figure 4.**
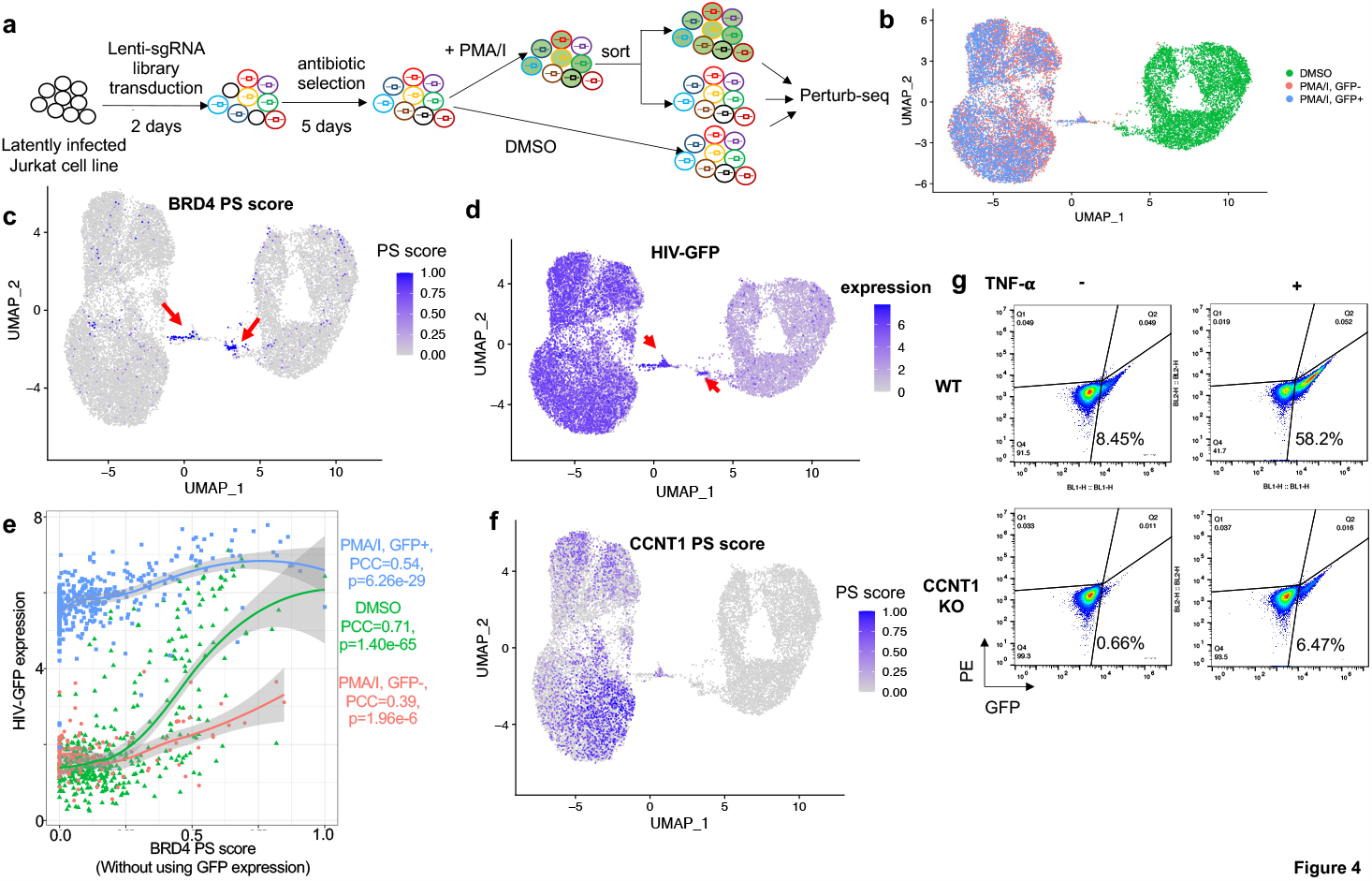
Perturb-seq on HIV latency. **a**, The experimental design of Perturb-seq. **b**, The UMAP plot of single-cell transcriptome profiles. Cells are colored by three different conditions. **c**, The distribution of BRD4 PS. **d**, The expression of HIV-GFP. **e**, The correlations between HIV-GFP expression and BRD4 PS that does not use HIV-GFP as target gene. **f**, The distribution of CCNT1 PS. **g**, The protein expression of HIV-GFP in response to CCNT1 knockout in different cell states (TNF-alpha vs non-stimulated).

We investigated gene functions using our PS framework. Among all perturbed genes, the PS of BRD4 (bromodomain containing 4) demonstrates a strong cell state-specific pattern, where a subset of cells with BRD4 perturbation has strong BRD4 PSs (named “BRD4-PS+ cells”) than other BRD4-perturbed cells (or “BRD4-PS-cells; **Fig. 4c**). BRD4-PS+ cells overexpress genes that are involved in known functions of BRD4^30,31^ including NF-kB/TNF-alpha signaling, hypoxia and apoptosis (**Fig. 4c-d, Supplementary Fig. S6b-d**). We examined whether the differences in BRD4 PS reflects the degree of BRD4 functional perturbation. We first checked the expressions of BRD4 “signature” genes from another published study^32^. Compared with BRD4-PS-cells, BRD4-PS+ cells have a much lower expressions of these signature genes (**Supplementary Fig. S6e**), indicating a stronger functional BRD4 perturbation. In addition, BRD4 has been shown to inhibit HIV transcription and activation in many studies, including our previous CRISPR screens^29,33^, consistent with the fact that HIV-GFP is one of the strongest up-regulated genes in BRD4-PS+ cells (**Supplementary Fig. S6f**). Furthermore, BRD4-PS+ cells have a stronger GFP expression (**Fig. 4d**) than other cells, confirming a stronger BRD4 functional perturbation in these cells.

To build a quantitative perturbation-expression relationship, we recalculated BRD4 PS without using HIV-GFP expression and examined how the scores are associated with a phenotype of interest (i.e., latent HIV-GFP expression) in different conditions (**Fig. 4e**). BRD4 PS correlation with HIV-GFP expression is cell-state dependent: in stimulated T cells (PMA/I treatment), a linear, positive correlation is observed regardless of the GFP expression. In contrast, a nonlinear relationship exists in unstimulated T cells (DMSO), where stronger BRD4 PS (>0.5) leads to a sharp increase in HIV-GFP expression (**Fig. 4e**).

Another gene, cyclin T1 (CCNT1), also displays heterogeneity in PS distribution: cells with CCNT1 perturbation have a high PS distribution only in stimulated cells (**Fig. 4f**). This is different from CCNT1 gene expression or guide distribution, which do not show such pattern differences between cell states (**Supplementary Fig. S7a**). Confirming our findings, the number of DEGs (cells with CCNT1 perturbation vs. cells expressing non-targeting guides) is over 100 in stimulated cells, but only 1 in non-stimulated cells (adjusted p value <0.001; **Supplementary Fig. S7b**). In particular, HIV-GFP is the strongest DEG in cells with CCNT1 perturbation, consistent with the known role of CCNT1 in activating HIV transcription.

CCNT1 is a key subunit of P-TEFb (positive transcription elongation factor b)/CDK9 complex that drives RNA transcription, including the transcription of HIV. The transcription elongation control of P-TEFb/CDK9 is a complicated process that is regulated by multiple mechanisms, including various T cell signaling pathways (e.g., NF-kB signaling), translation control, and epigenetic modification (reviewed in ^34^). The activities of these factors are different in different states of T cells (e.g., NF-kB; **Supplementary Fig. S7c**), which may explain the differences of CCNT1 PSs. Despite the strong cell state dependency of CCNT1 PS, PS shows weak correlation with HIV-GFP within one cell state (**Supplementary Fig. S7d**), which is different from BRD4 PS (**Fig. 4e**).

To further confirm our finding that different cellular states affect the transcriptomic responses of CCNT1 perturbation, we stimulated Jurkat cells using a different agonist (TNF-alpha). To measure the downstream effect of CCNT1 perturbation, we sorted cells by expression of HIV-GFP, which is the strongest down-regulated gene upon CCNT1 knockout (**Supplementary Fig. S7b**), and whose expression is known to be regulated by CCNT1^35,36^. Indeed, with the presence of TNF-alpha, CCNT1 knockout leads to a strong reduction in HIV-GFP expression (over 50% reduction), while such reduction is much smaller (<5% reduction) in cellular states without TNF-alpha stimulation (**Fig. 4g**).

Collectively, these results demonstrated that PS is a powerful computational framework for investigating cofactors (cell states, other genes) that drive transcriptomic responses upon gene perturbation.

### PS enables identification of novel cell-type dependent gene functions in regulating pancreatic cell differentiation from multiplex single-cell transcriptomics

Besides Perturb-seq, multiplexing cells with different perturbations are also used to measure single-cell responses to perturbation^2,19^. A mixture of cells from different perturbations can be sequenced at the same time, and the identity of cells can be established using various methods including cell hashing^37^, the expressions of pre-defined barcodes^38^, or a combination of random barcodes^39^. We therefore tested our PS framework on pooled single-cell transcriptomics of different perturbations to study the functions of lineage regulators during human pancreatic differentiation. By using an established in vitro human embryonic stem cell (hESC) pancreatic differentiation system, we generated cells corresponding to early stage (definitive endoderm, DE) and middle stage (pancreatic progenitor, PP) pancreas development. To test the performance of PS framework and uncover the functions of unknown regulators, we picked ten clonal hESC lines with the homozygous knockout of four genes (**Supplementary Table S4**), including two known pancreatic lineage regulators (HHEX, FOXA1) and two uncharacterized candidate regulators from previous genetic screens (OTUD5, CCDC6)^40,41^. These clones are then labelled with different LARRY (Lineage and RNA recovery) DNA barcodes^38^, pooled together and differentiated into DE and PP stages using established protocols^40^. Finally, the single-cell expressions of these cells were profiled via 10X genomics Chromium platform (**Fig. 5a**). The clone information of each cell was identified from LARRY barcodes. Among 26,286 single cells that passed the quality control measurements, over 97% (25,694/26,286) of the cells had at least one barcode detected, and over 80% (20,678/25,694) were identified as singlets and retained for downstream analysis. UMAP clustering revealed different known cell types during pancreatic differentiation, based on the expression markers of known cell types (**Fig. 5b; Supplementary Fig. S8**), including DE, PP, liver/duodenum progenitor (LV/DUO), endocrine precursor (EP), and cells in transition stages (e.g., DE in transition, PP in transition).

**Figure 5.**
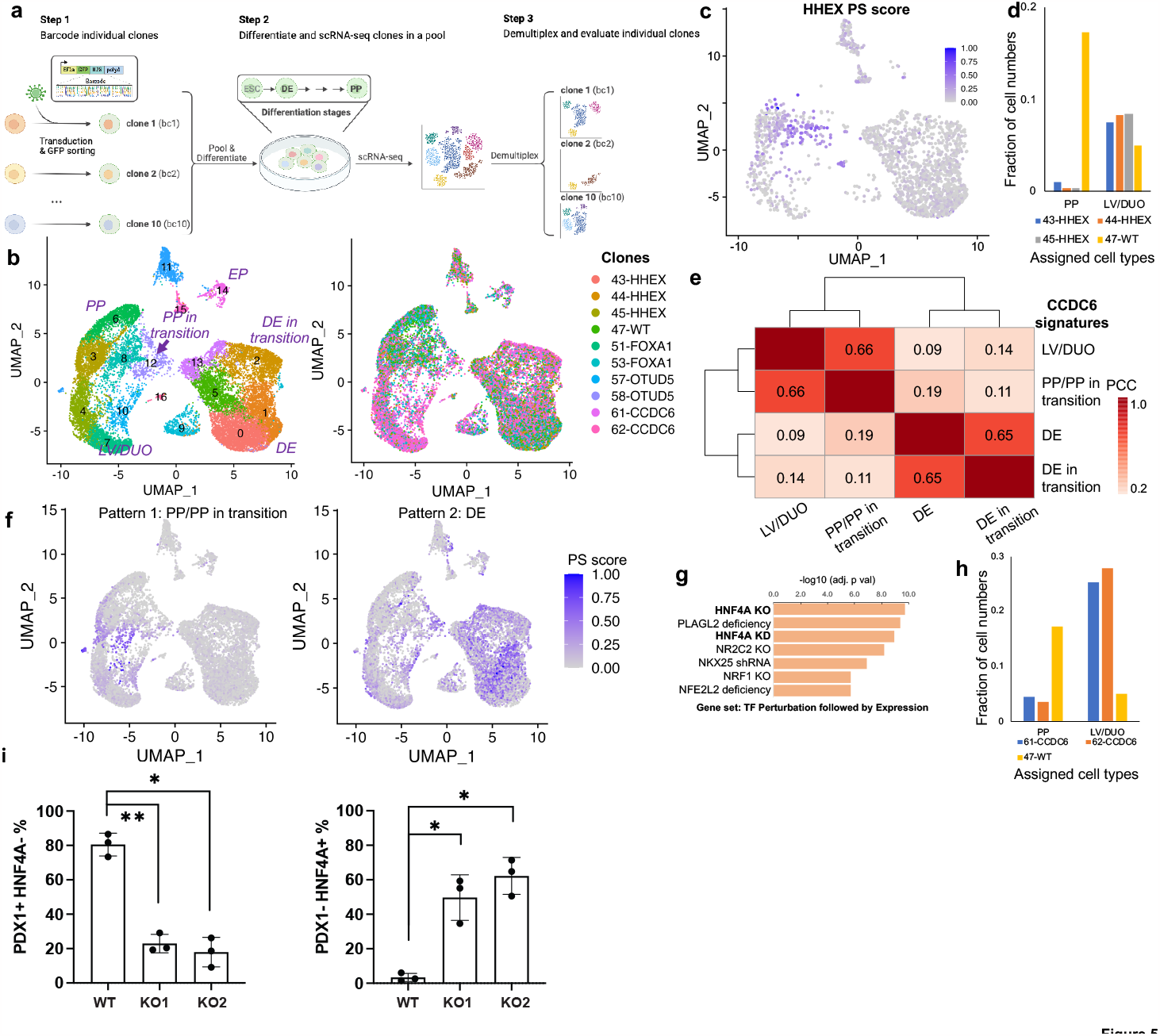
Pooled scRNA-seq on pancreatic differentiation. **a**, Experimental design of multiplexing scRNA-seq on the knockout clones of different genes. **b**, The UMAP plot of single-cell transcriptome profiles, colored by different clusters (left) or clones (right). **c**, The PS distribution of HHEX. **d**, The percentage of cells in PP/LV/DUO cell types from different clones. **e**, The correlations of CCDC6 PSs calculated from different HHEX cell types. The Pearson Correlation Coefficient (PCC) is calculated from all cells with CCDC6 knockouts and is shown as numbers on the heatmap. **f**, Two different distribution patterns of CCDC6 PSs. **g**, The top enriched GO terms of DEGs from PP/PP in transition. Enrichr was used to perform enrichment analysis. **h**, The percentage of cells in PP/LV/DUO cell types from CCDC6 clones. **i**, The percentage of cells with PDX1+ (a PP marker) or HNF4A+ (a LV marker) by flow cytometry sorting. The data is based on two CCDC6 knockouts (KO1, KO2) and one wild-type (WT) control. Three independent replicates are performed for each condition. The multiple comparison-adjusted p value is calculated by one-way ANOVA test. *p<0.05, **p<0.01.

We next applied the PS framework to the pooled single-cell RNA-seq datasets containing different knockout clones. Among all knockout genes, HHEX PS is high in cells whose type is between two different differentiated cell types (PP and LP/DP; **Fig. 5c; Supplementary Fig. S8**), consistent with the known function of HHEX as a key determinant of cell fate decision, whose deletion drives DE cell differentiation towards LP/DP, rather than PP^40^. Indeed, *HHEX* knockout led to a much fewer percentage of cells that are annotated as PP (**Fig. 5d**). The PS of FOXA1, another key transcription factor during PP differentiation, is strong in DE and PP cell types, consistent with the specific expression pattern of FOXA1 in DE/PP cell types (**Supplementary Fig. S9a-c)**.

As in our previous genome-wide CRISPR screens, CCDC6 is one of the top hits whose perturbation hinders PP differentiation^40,42^. However, the exact function of CCDC6 during pancreatic differentiation is largely unknown. CCDC6 may have different functions at different cell types, evidenced by the few overlaps of DEGs between different cell types (**Supplementary Fig. S9d-f**). To investigate these different functions, we calculated PSs from the DEGs from four major cell types in the dataset (DE in transition, DE, PP/PP in transition, and LV/DUO). An unbiased clustering on these CCDC6 PSs demonstrated two distinct distributions across cell types (**Fig. 5e**), where scores calculated from late-stage cell types including PP/PP in transition/LV/DUO (“pattern 1”) are distributed differently from scores calculated from early-stage cell types including DE in transition/DE (“pattern 2”; **Fig. 5f; Supplementary Fig. S10a-b**), implying different behaviors of CCDC6 perturbation at different cell types. Indeed, functional analysis on DEGs leading to both patterns have distinct enrichment terms. In early-stage cell types, DEG genes are enriched in the targets of stem cell transcription factors (e.g., SOX2, POU5F1, NANOG) and cell cycle regulation (**Supplementary Fig. S10c-e**), consistent with the known function of *CCDC6* as a cell cycle regulator^43,44^. In contrast, DEGs in late-stage cell types are primarily the targets of HNF4A, a key transcription factor that drives LP/DP differentiation (**Fig. 5g; Supplementary Fig. S10f**). The expressions of these transcription factors (SOX2, HNF4A) are among the up-regulated genes in both programs, respectively (**Supplementary Fig. S9d-e**). Furthermore, compared with wild-type cells, *CCDC6* knockout cells have a much lower percentage of PP cells and a higher percentage of LP/DP cells (**Fig. 5h**). Collectively, these results imply that *CCDC6* has different functions for early vs. late-stage cell types. Especially in late-stage cell types, *CCDC6* knockout drives cell differentiation towards LV/DUO cell types rather than PP cell types.

To further validate the prediction results of CCDC6, we performed flow cytometry analysis to evaluate the effects of *CCDC6* knockout on the composition of late-stage cell types (PP/LV/DUO). We examined the percentage of HNF4A+ cells, a marker for LV population, and PDX1+ cells, a marker for PP population. Indeed, both clones of *CCDC6* knockout greatly reduced PDX1+ population and increased HNF4A+ population in three biological replicates (**Fig. 5i; Supplementary Fig. S11**), confirming our finding on the enrichment of CCDC6 PS in LP/DP populations (**Fig. 5f-g**).

## Discussion

Understanding cellular responses to perturbations is a central task in modern biology, from studying tumor heterogeneity to developing personalized medicine. These perturbations may be genetic (e.g., knocking out genes or non-coding elements), chemical (e.g., drug treatments), mechanical (e.g., pressure) or environmental (e.g., temperature changes). Single-cell genomics profiles of perturbations are commonly used to investigate the mechanisms of perturbations. Many technologies, including Perturb-seq and sci-Plex, provide a high-content readout of the results of systematically perturbing many genes or non-coding elements. Despite rapid technological advancements, a major bottleneck is the lack of a computational model to fully unlock the potential of high-content perturbation, especially for discovering novel biological insights from the data. Here we introduce the PS framework to model the heterogenous transcriptomic responses of perturbations and to enable novel biological discovery from modeling perturbation heterogeneity.

Partial gene perturbation is common in perturbation experiments. Partial perturbations may come from dose-controlled drug treatment, gene editing technology that does not fully knockout gene function (e.g., RNA-or CRISPR-interference, epigenome editors), or from CRISPR/Cas9 that generates random DNA editing outcomes. We demonstrated the outstanding performance of our PS method over existing methods in quantifying partial gene perturbation. Specifically, partial perturbation identification enables the analysis of dose-dependent effect, which is demonstrated in this study using various datasets.

More importantly, PS enables novel biological investigations, including analysis of perturbation dosage without the need to titrate perturbation strength and identification of cell-intrinsic and extrinsic biological factors that regulate perturbation responses. In the latter case, the PS, ranging between 0 and 1, no longer represents the quantity of partial perturbation, but instead represents the strength of the perturbation outcome. Therefore, PS becomes a convenient tool to identify cell context that determines perturbation outcome. We demonstrated the application of PS in various biological problems, including T cell activation, essential gene function, latent HIV-1 virus expression, and pancreatic cell differentiation. Importantly, our PS model leads the discovery of novel CCDC6 functions that are cell type dependent, whose role as a regulator during pancreatic and liver cell fate decision is experimentally validated.

Partial perturbations of gene functions contribute to the complexity of many biological processes. For example, “haploinsufficient” genes are able to cause disease phenotypes when 50% of their functions are disrupted, while “haplosufficient” genes will require a nearly complete gene knockout. However, we currently lack a method to investigate the phenotypes of partial gene perturbations or to efficiently perform dosage analysis at a large scale. Current approaches, such as introducing mismatches to guide RNAs to modulate the effects of CRISPRi^21^ or Cas13^28^, require a complex design of a specific CRISPR system. Here we demonstrated that both CRISPR knockout (e.g., **Fig. 2f, Fig. 4e**) and CRISPRi naturally introduce partial perturbation effects, which can be used to study the dose effect of partial gene perturbations on downstream gene expressions or a phenotype of interest. Our PS framework is versatile, enabling the dosage analysis using various perturbation methods (e.g., CRISPRi or CRISPR knockout) and assays (e.g., Perturb-seq or multiplex scRNA-seq).

Results from genetic perturbations (e.g., via CRISPR/Cas9) are informative for drug development, and confirmations from genetic perturbation experiments are usually required to demonstrate the feasibility of candidate drug targets. However, titrating pharmaceutical interventions are easy (e.g., by using different doses of drugs), while it is much more difficult to precisely control the degree of genetic perturbations. Our PS framework provides a convenient alternative to dose-dependent perturbations, especially genetic perturbations, and their associations with phenotypic changes, which will be informative in designing drugs. For example, BRD4 is the primary target of bromodomain inhibitors (BETi), many of which have been proposed as candidates of latency reversing agents (LRAs) to reactivate latent HIV-1 expression. The distribution of BRD4 PSs (**Fig. 3**) reveals that stronger perturbation effects are needed to induce the desired phenotype, in this case, the expression of HIV-GFP (**Fig. 3**). Since BRD4 is an essential gene, a strong BRD4 perturbation may lead to unexpected toxicity, thereby limiting the efficacy of BETi. Indeed, our previous study^29^ demonstrated that 10-1000x higher doses of JQ1, a commonly used BETi, are needed to induce latent HIV-1 expression at a similar level with other potent LRAs. Our results further warrant the development of synergistic drug combinations to mitigate the narrow therapeutic window of BETi, which is currently tested in many studies.

Our PS analysis provides a general framework to analyze several major sources that contribute to the heterogeneity of perturbation responses: the strength of perturbation *per se* (e.g., **Fig. 1i, 3d;** BRD4 in **Fig. 4c**), compensations to perturbation especially on essential genes (e.g., proteosomes; **Fig. 3g**), and cell type/state specificity (e.g., T cell states in **Fig. 4**; differentiation cell types in **Fig. 5**). Importantly, cell type/state is linked to perturbation responses in three distinct ways: cell type/state may change as a result of perturbation (e.g., CCDC6 and HHEX in Fig. 5); cell type/state serving a critical context to define perturbation responses (e.g., T cell states in response to CCNT1 perturbation in **Fig. 4f-g**); and cell type/state as a confounding factor that drives perturbation responses (e.g., BRD4 perturbation heterogeneity in unstimulated T cells in **Fig. 4c**). Compared with other methods, PS is currently the only method to analyze heterogeneity of perturbation responses from all these aspects.

Confounding factors are the major sources of variation when analyzing single-cell perturbation effects. These confounding factors can be modeled explicitly (e.g., using generalized linear models) if confounding source is known; or be detected and corrected using mathematical or statistical approaches including matrix factorization (e.g., using GSFA^45^) or independent component analysis (e.g., using CINEMA-OT^46^). In contrast, PS does not explicitly model confounding factors. Instead, PS scores can be used in combination with methods that remove confounding sources of variation, or to detect these confounding factors that contribute to the heterogeneity in perturbation responses (e.g., **Fig. 4c**).

Importantly, many confounding factors defined in previous methods^16,46^ are not always confounding; instead, they can be used to discover novel biological insights, as are shown in this study (e.g., perturbation efficiency, cell type/state). The orthogonal algorithmic design of PS compared with existing methods also allows the combination of PS with these methods to simultaneously remove confounding factors and measure the strength of perturbation responses.

One limitation of PS is its power in detecting drastic changes in cell types or states. For example, even moderate perturbations on essential gene functions affect cellular viability^47,48^. In this case, single-cell profiling only captures surviving cells that are resistant to essential gene perturbations in various mechanisms (e.g., expression compensation in **Fig. 3**), and largely misses dead cells due to essential gene dysfunction. Consequently, due to this “survival bias”, PS probably only reflects the perturbation responses in a fraction of cells, rather than the full spectrum of perturbations. To overcome this limitation, PS can combine with recently developed prediction methods that predict the responses of perturbations, even if cells between perturbed/non-perturbed states are unevenly distributed^49^.

## Methods

### The Perturbation-response Score (PS) framework

Estimating PS proceeds in three steps, as illustrated in Figure 1c: target gene identification (Step 1), average perturbation effect estimation using a previously published scMAGeCK (Step 2), and PS estimation using constrained optimization (Step 3).

#### Step 1: target gene identification

We first performed differential expression analysis between cells with certain perturbation (e.g., knocking out gene *X*) and negative control cells. In most cases, negative control cells are cells that express non-targeting guide RNAs (in Perturb-seq), or wild-type cells (in pooled scRNA-seq). In Perturb-seq with high MOI condition, these cells may come from cells that do not have a particular perturbation. We used Wilcoxon rank sum test (implemented in Seurat) to identify and rank differentially expressed genes. Top genes were then selected as potential target genes of the specific perturbation. The maximum and minimum numbers of top genes can be specified by the user. Alternatively, users can provide the list of target genes for each perturbation, based on prior knowledge, therefore skipping the differential expression analysis in this step.

#### Step 2: average perturbation effect estimation

We used the linear regression module in scMAGeCK (scMAGeCK-LR) to estimate the average perturbation effect. scMAGeCK-LR takes the expressions of all target genes (identified in Step 1) in all cells as input and outputs a β score, which is conceptually similar to log fold change. There are two advantages of using β score, instead of simply using the log fold changes in Step 1. First, scMAGeCK-LR naturally supports datasets from high MOI Perturb-seq, where one cell may express multiple guides targeting different genes. Second, scMAGeCK-LR is able to estimate average perturbation effects of multiple perturbations (e.g., genome-scale perturbations) in one step, while a naïve DEG analysis can only calculate LFC for each perturbation.

The mathematical model of scMAGeCK-LR is described as follows. Let *Y* be the log-transformed, *M*N* expression matrix of *M* single cells and *N* target genes. These genes are the union of all target genes for all *K* perturbations, extracted from Step 1. Let *D* be the *M*K* binary cell identity matrix of *M* single cells and *K* perturbations, where *d*_*jX*_ = 1 if single cell *j* contains sgRNAs targeting gene *X* (*j* = 1,2, …, *M*; *X* = 1,2, …, *K*), and *d*_*jX*_ = 0 otherwise. *D* can be obtained from the detected guide RNA expression matrix from Perturb-seq or from the prior sample information from pooled scRNA-seq. The effect of target gene knockout on all expressed genes is indicated as a β score in a matrix B with size *K*N*, where *β*_*XA*_ > 0 (< 0) indicates gene *X* is positively (or negatively) selected on gene *A* expression, respectively. In other words, gene *X* knockout increases (or decreases) gene *A* expression if *β*_*XA*_ > 0 (< 0), respectively.

The log-transformed expression matrix *Y* is modeled as follows:

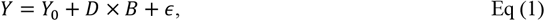

where *Y*_0_ is the basal expression level of all genes in an unperturbed state, and *ϵ* is a noise term following a Gaussian distribution with zero means. *Y*_0_ can be estimated from negative control cells (e.g., wild-type cells or cells expressing non-targeting guides), or be modeled using the expressions of neighboring negative control cells (e.g., the approach used by mixscape^16^). The value of B can be estimated using ridge regression:

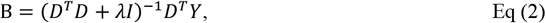

where *I* is the identity matrix, and *λ* is a small positive value (default 0.01).

#### Step 3: PS estimation using constrained optimization

We revise Eq (1) to incorporate PS. Here, the log-transformed expression matrix Y is modelled as follows:

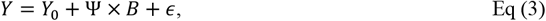

Where Ψ is the non-negative, raw PS matrix with the same size as *D* in Step 2 (*M*K)*. Each element ψ_jX_ in Ψ indicates the raw PS of cell *j* of perturbing gene *X*. Here, *B* is the β score matrix which is estimated in Step 2. We find the value of Ψ to minimize the squared error of predicted and observed expressions of all genes within all cells, subject to constraints and regularization terms:

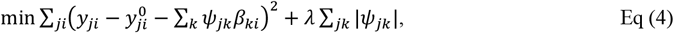

subject to the following constraints:

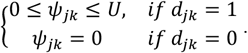

Here, *U* is a positive value indicating the upper bound of raw Ψ values, and *d*_*ik*_ is the value of the binary cell identity matrix in Step 2. 1 ≤ *j* ≤ *M* is the index of single cells, 1 ≤ *i* ≤ *N* is the index of target genes, and 1 ≤ *k* ≤ *K* is the index of perturbations.

Because we are imposing non-negative constraints to Ψ, the absolute operator can be removed from the objective function in Eq (4) and can be rewritten as

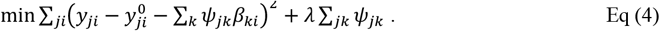

This becomes a constrained quadratic optimization problem where the best solution can be easily achieved using methods like Newton’s method. The final, normalized PS is to scale values of *ψ*_*ik*_ to [0,1]:

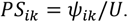

We implemented this framework as part of the scMAGeCK pipeline^18^. The PS source code, documentation and tutorials can be found on Github: https://github.com/davidliwei/PS

### Simulated datasets

The eight simulated datasets are generated by the simulator scDesign3^46^ with modifications for Perturb-seq. The simulation utilizes scDesign3’s parametric model to capture the characteristics of the user-inputted reference data, specify the desired ground truth, and simulate synthetic cells via sampling from the model (to be detailed in **Steps 1-4** below). The reference data is the real scRNA-seq dataset with the gene Nelfb perturbed in some mouse T cells^47^; the cells with Nelfb perturbed are referred to as *knockout cells*, and the cells with Nelfb unperturbed serve as the negative control and are referred to as *wild-type cells*. Based on the same reference data, the eight simulated datasets are generated under eight different settings. Each setting corresponds to a combination of two simulation parameters’ values: the number of Nelfb’s downstream genes (i.e., the genes whose expression levels are affected by Nelfb’s knockout; with candidate values 0, 10, 200, and 500) and the perturbation efficiency (with candidate values 50% and 100%). The candidate downstream genes of Nelfb are the top differentially expressed (DE) genes identified from the bulk RNA-seq data of the same biological sample (from the second sheet in the Excel file from Wu et al.’s Supplementary Data 1^48^). Thus, we have 4×2=8 simulated datasets in total.

Before running the simulation, we pre-process the scRNA-seq dataset and the bulk DE gene rank list.

1. First, we perform the same quality control as in the dataset’s original publication^49^. Specifically, cells are retained only if their numbers of detected genes are between 1,000 and 5,000, and their UMI counts have less than 12% mitochondrial counts.
2. Second, we impute and amplify the gene-by-cell count matrix of the wild-type mouse cells to enhance the perturbation effects in the simulated data. Specifically, we first impute the wild-type count matrix using scImpute^50^ (default version 0.0.9) to reduce the sparsity. Then we multiply the imputed count matrix by an amplification factor of 10 to increase the range of gene expression levels.
3. Third, we construct a gene-by-cell count matrix by combining the wild-type cells in the post-imputation- and-amplification wild-type count matrix and the knockout cells in the knockout count matrix. By the end of this step, the dimension of this combined matrix is (*P*+1)×*N*, with rows corresponding to *P*+1 genes (Nelfb and *P* other genes) and columns corresponding to *N* cells, which consist of *N*^wt^ wild-type cells and *N*^ko^ knockout cells.
4. Fourth, we extract the row corresponding to Nelfb as a vector, which contains Nelfb’s counts in all cells (an *N*-dimensional vector denoted as *C*, where *C*_*j*_ is Nelfb’s count in cell *j*), and we denote the remaining *P* rows as a *P*×*N* matrix **Y**, where *Y*_*ij*_ is gene *i*’s count in cell *j*.
5. Fifth, using **Y**, we refine the list of bulk DE genes by excluding the DE genes that correspond to zero rows in **Y** or do not correspond to any rows in **Y**.
6. Lastly, to reduce the computation time for data simulation, we use the scran package^51^ to select 3,000 highly variable genes in **Y**. We only keep the union of these 3,000 highly variable genes and the refined bulk DE genes as the rows in **Y**. The number of the kept genes is 3,390, so the dimension of **Y** is 3,390×*N*.

Additionally, we know which cells have Nelfb perturbed; thus, we have another *N*-dimensional binary vector denoted as *K*, where *K*_*j*_ indicates whether the *j*-th cell has Nelfb perturbed or not; that is, *K*_*j*_ = 0 means the *j*-th cell is a wild-type cell, and *K*_*j*_ = 1 means the *j*-th cell is a knockout cell. *K* and *C* are used as two covariate vectors, and **Y** is used as the reference count matrix for scDesign3. Finally, we modify scDesign3 by using **Y**, *C, K*, the refined DE genes, the number of Nelfb’s downstream genes, and the perturbation efficiency to simulate data in the following four steps:

#### Step 1: modeling each gene’s marginal distribution independently

For each gene *i*, if it is a downstream gene of Nelfb, we assume that *Y*_*ij*_, conditional on *C*_*j*_, follows a zero-inflated negative binomial (ZINB) distribution with the mean parameter *μ*_*ij*_, the dispersion parameter *ϕ*_*i*_, and the zero-inflation probability parameter *v*_*ij*_. Otherwise, if gene *i* is not a downstream gene of Nelfb, we assume that *Y*_*ij*_ follows a ZINB distribution with the mean parameter *μ*_*i*_, the dispersion parameter *ϕ*_*i*_, and the zero-inflation probability parameter *v*_*i*_. This marginal distribution for each gene is specified by a generalized additive model for location, scale, and shape (GAMLSS). Without loss of generality, we define the first *D* genes in **Y** to be the top *D* DE genes in the refined DE gene list (*D* ∈{0, 10, 200, 500}); we treat these top *D* DE genes as the *D* downstream genes of Nelfb. Then we modify scDesign3’s original code implementation so Nelfb’s downstream genes and non-downstream genes have different marginal distributions: a downstream gene’s marginal distribution in each cell *j* depends on *C*_*j*_, Nelfb’s count in cell *j*; a non-downstream gene’s marginal distribution in each cell *j* is irrelevant to *C*_*j*_.

For Nelfb’s downstream gene *i* = 1, …, *D*:

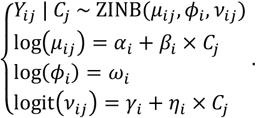

For Nelfb’s non-downstream gene *i* = *D* + 1, …, *P*:

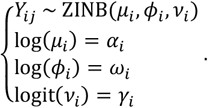

After parameter estimation by the R package gamlss, the fitted distribution of *Y*_*ij*_ | *C*_*j*_, for *i* = 1, …, *D*, is denoted as 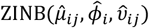 with the 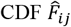; the fitted distribution of *Y*_*ij*_, for *i* = *D* + 1, …, *P*, is denoted as 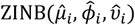 with the 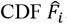. The other parameters including *α*_*i*_, *β*_*i*_, *γ*_*i*_, and *η*_*i*_ are estimated as 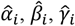, and 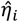 for each *i* respectively.

#### Step 2: modeling genes’ joint distribution using the Gaussian copula

To approximate the pairwise gene-gene correlations in the reference dataset, scDesign3 utilizes a multivariate statistical technique, the Gaussian copula. Given each gene’s marginal distribution fitted in Step 1, scDesign3 approximates the multivariate joint distribution of the *P* genes in cell *j* as

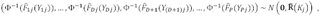

where Φ^−1^(⋅) denotes the inverse of the cumulative distribution function (CDF) of the standard Gaussian distribution, **0** is the *P*-dimensional zero vector, and 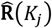 is the estimated *P* × *P* gene-gene correlation matrix of the Gaussian copula conditional on the value of *K*_*j*_. Specifically, since *K*_*j*_ is binary, we have two estimated gene-gene correlation matrices, one for the wild-type cells (*K*_*j*_ = 0) and the other for the knockout cells (*K*_*j*_ = 1). For 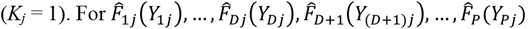, a technique called “distributional transform” is used to make the CDFs continuous; see Sun et al.^52^ for a detailed explanation.

#### Step 3: modifying the fitted parameters

Since we want to generate synthetic datasets with two perturbation efficiencies, for each downstream gene *i* = 1, …, *D*, we modify the mean parameters for all downstream genes in the knockout cells to reflect the user-specified perturbation efficiency. Without loss of generality, we assume the first 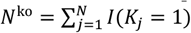 of the *N* cells as the knockout cells. Then, we update the mean parameters for Nelfb’s *D* downstream genes in the *N*^ko^ knockout cells (i.e., 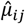 for *i* ∈{1, …, *D*}, *j* ∈{1, …, *N*^ko^}) based on the user-specified perturbation efficiency as follows.

For the 50% perturbation efficiency: We randomly sample *N*^ko^ values from {*C*_*j*_, *j* ∈{*N*^ko^ + 1, …, *N*}} (i.e., Nelfb’s counts in the wild-type cells) and multiply the sampled *C*_*j*_ values by 0.5 to represent the 50% perturbation efficiency. We store these sampled and scaled values by 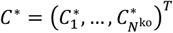 as Nelfb’s counts in the *N*^ko^ synthetic knockout cells to be simulated. Then, we modify the mean parameters for the *D* downstream genes in the *N*^ko^ synthetic knockout cells (for *i* ∈{1, …, *D*}, *j* ∈{1, …, *N*^ko^}) as

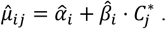

For the 100% perturbation efficiency: *C*^∗^ becomes a zero vector with length *N*^ko^, and we modify 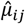 for *i* ∈{1, …, *D*}, *j* ∈{1, …, *N*^ko^} in the same way as above.

We do not change any estimated mean parameters for the *D* downstream genes in the *N*^wt^ wild-type cells, any estimated mean parameters for the non-downstream (*P* − *D*) genes in all *N* cells, any estimated dispersion parameters, or any estimated zero-inflation probability parameters.

Moreover, we use *S* to denote an *N*-dimensional vector representing Nelfb’s counts in the *N* synthetic cells, with the counts in the first *N*^ko^ synthetic knockout cells set above based on the perturbation efficiency, and the counts in the last *N*^wt^ synthetic wild-type cells same as those in the real *N*^wt^ wild-type cells. That is, 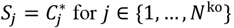for *j* ∈{1, …, *N*^ko^}, and *S*_*j*_ = *C*_*j*_ for *j* ∈{*N*^ko^ + 1, …, *N*}.

#### Step 4: generating synthetic data with the fitted model and modified parameters

First, we independently sample *N*^wt^ Gaussian vectors of length *P* from the estimated *P*-dimensional multivariate Gaussian distribution 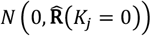 and *N*^ko^ Gaussian vectors of length *P* from the estimated *P*-dimensional multivariate Gaussian distribution 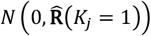. Together, we stack these *N* = *N*^wt^ + *N*^ko^ vectors 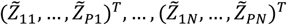 by row into a *P*×*N* Gaussian matrix 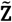.

Given the parameter est*i*mates (mod*i*f*i*ed or not) from Step 3, we convert the *P*×*N* Gauss*i*an matr*i*x 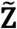 into a *P*×*N* ZINB count matr*i*x 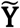 as

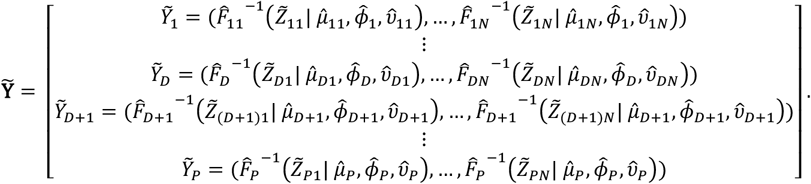

Lastly, we combine 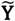 w*i*th *S* by row into a (*P+*1)×*N* matrix, obtaining the final (*P*+1)×*N* synthetic count matrix 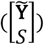.

### Genome-scale Perturb-seq on Jurkat cells

#### Perturb-seq

We performed genome-scale Perturb-seq on Jurkat E6 cell line expressing dCas9-KRAB as our model to study. We transduced them with a genome-wide CRISPRi CROP-seq library at a high MOI. After infection, we split the cells into two populations, including untreated cells and activated cells (cells treated with anti-TCR and anti-CD28 antibodies for approximately 24 hours to stimulate TCR signaling). Cells were then labelled with cell hashing antibodies. Multiple labels were used for the activated population to help with cell multiplet detection. Cells were loaded on 16 channels of a 10x Chromium X instrument. We loaded 115 000 cells per channel, and the expected recovery rate was 60 000 cells per channel, including 24% multiples. Samples were pooled unequally before they were loaded on the ChromiumX: 10% untreated cells, 90% treated cells. A sequencing library was prepared using 3’Chemistry with a targeted primer panel: custom multiplex PCR step to enrich for specific transcripts. Libraries were sequenced on NovaSeq S4 PE100 in asymetric read mode (R1: 28 cycles; R2: 172 cycles), with PhiX concentration of 1%. The expected coverage is around 9 000 ∼ 10 000 input reads per cell.

#### Hash oligos

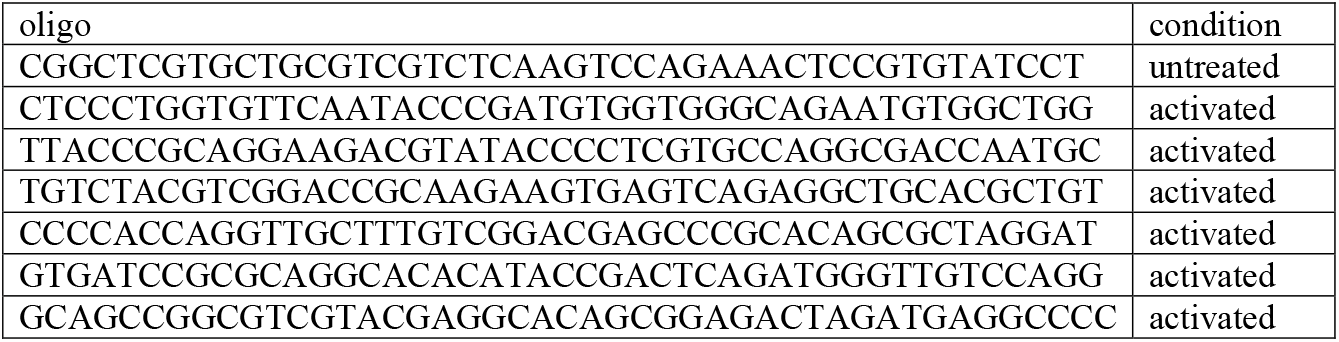

#### sgRNA library design

The genome-wide CRISPRi sgRNA library was designed to target the transcription start site (TSS) coordinates, calculated from publicly available FANTOM CAGE peaks data. In total, 18 595 genes were targeted, with 4 sgRNAs per gene. On top of that, we designed another CRISPRi library targeting 3220 genes with 4 sgRNAs per gene. This library was designed using Jurkat-specific TSS, which were calculated from public Jurkat CAGE-seq datasets. Both libraries were combined into a final library targeting 3220 genes with 8 sgRNA/gene and 15 375 genes with 4 sgRNA/gene.

#### Targeted primer panel

The primer panel for targeted transcriptomic readout consisted of 374 target genes from several categories:

| Source | Number of targets (genes) |
| --- | --- |
| bulk RNA-seq DEG, 50 top up/downregulated | 100 |
| T-cell related genes | 51 |
| KEGG TCR signaling pathway | 35 |
| RNA-binding proteins | 27 |
| Replogle*: unfolded protein response | 51 |
| Replogle: proteasome | 41 |
| Replogle: NFkB | 20 |
| Replogle: cell cycle | 29 |
| Replogle: targets of nonsense-mediated decay | 5 |
| Controls (cell cycle, mitochondrial, Cas9) | 15 |
| Total number of targets | 374 |

\* “Replogle” prefix means that this category of targets was derived from the published genome-scale Perturb-seq dataset<sup>50</sup>.

#### Data preprocessing

For each of the 16 channels, all 3 kinds of sequencing libraries (mRNA, sgRNA, cell hashing) were indexed using the same Illumina index sequence. We obtained high-quality scRNA-seq data of over 586,000 single cells after quality control, with a median 13 guides detected per cell. We obtained an average of 400 cells per gene perturbation. STAR and STAR solo 2.7.10a were used to map transcriptomic reads against a custom gtf annotation, which was based on gencode.v34.annotation.gtf (hg38). Reads that did not map to the transcriptome reference were then mapped & counted using STAR solo against a custom fasta reference with guide sequences and a fasta reference with hash label sequences. STAR Solo output transcriptome matrices were first filtered using an approach similar to 10x cellranger EmptyDrops filtering, which retained cells with at least 10 % UMI count of the 99^th^ percentile UMI counts of the top expected cells number. Then, an initial Seurat object was created from those filtered transcriptome matrices using the CreateSeuratObject function with following parameters: min.cells = 5; min.features = 10; all other parameters at default values. Outlier cells were filtered out by mithochondrial and mRNA content (percent.mt, nCount_RNA). In order to detect cell multiplets and determine cell population (untreated or activated), cell labels (also known as hashes) were called using the MULTI-seq approach (deMULTIplex::classifyCells in R). Only cells with exactly 1 known label were kept. Then, sgRNA calling was conducted using a binomial test, with total sgRNA UMI counts used to derive background frequencies. A threshold of 0.05 on Benjamini-Hochberg corrected p-values (per channel) was use to generate the final calls. The sgRNA assays are sparse matrices containing 1, where the respective cell is considered to be carrying the respective sgRNA and 0 elsewhere. Following that, all results from steps above from 16 channels were merged together, and merged counts were normalized using NormalizeData and scaled using ScaleData. Cell cycle scoring was performed via CellCycleScoring, PCA was calculated using RunPCA, and UMAP was calculated on first 30 principal components using RunUMAP in Seurat.

### HIV latency Perturb-seq

We used a previously established cell line model of HIV latency^40^. In this model, Jurkat cells were infected with an HIV vector with GFP tied to the LTR promoter, resulting in a positive GFP signal as a measurement of viral transcription reactivation and HIV latency reversal. These cells, which already express Cas9, were transduced with a lenti-sgRNA library. The lenti-sgRNA library (MilliporeSigma; LV14, U6-gRNA-10x:EF1a-Puro-2a-BFP) was designed to target 10 genes, with 3 gRNAs per gene. In addition to non-targeting controls, the library contained five positive regulators (NFKB1, CCNT1, PRKCA, TLR1, MAP3K14) and five negative regulators (NFKBIA, NELFE, HDAC2, BRD4, BIRC2) of HIV transcription. Transduction was carried out on 850,000 cells at an MOI of 0.3 using 8ug/ml polybrene in 2 ml of RPMI containing 10% FBS and 1% penicillin–streptomycin. The media was replaced 24 hours later with fresh media without polybrene. Two days after transduction, the cells were selected for using 1.5 ug/ml puromycin for 5 days. After selection, the cells were split evenly into three groups. One-third of the cells were kept in culture with no drug added, and two-thirds of the cells were stimulated with PMA/I (50ng/ml PMA in combination with 1 μM Ionomycin). After 16 hours, the stimulated cells were sorted into GFP+ and GFP-populations. All three samples were then analyzed following the 10x Genomics single-cell sequencing protocol. Sequencing data, encompassing gene expression and CRISPR guide capture libraries, underwent demultiplexing and processing using Cell Ranger (version 6.1.2). The resulting feature-barcode matrices from three samples were then merged, and subsequent analysis was carried out utilizing the Seurat R package (version 4.3.1). To ensure data quality, cells were excluded if the number of expressed genes was greater than 7,500 or fewer than 200. Additionally, cells were removed if the percentage of mitochondrial reads exceeded 15%. Single cells harboring more than one detected sgRNA sequence, attributable to either multiple sgRNA transductions or the presence of multiple cells in a single-cell droplet, were also excluded from the analysis. Following quality control measures, merged counts underwent normalization and scaling. PCA was computed based on the top 2,000 highly variable genes. Subsequently, clustering and UMAP embeddings were performed using default parameters. To gain further insights into the biological significance of the obtained clusters, enrichment analysis was conducted utilizing Enrichr (PMID: 27141961).

### Pancreatic differentiation clones and pooled single-cell RNA-seq

#### Culture of hESC

Generation of KO hESCs was described in published studies, including HHEX KO H1 and HUES8 cell lines^53^, FOXA1 KO HUES8 cell lines^54^, OTUD5 KO HUES8 cell lines, and CCDC6 KO H1 cell lines^41^. Cells were regularly confirmed to be mycoplasma-free by the Memorial Sloan Kettering Cancer Center (MSKCC) Antibody & Bioresource Core Facility. KO and WT hESCs were maintained in Essential 8 (E8) medium (Thermo Fisher Scientific, A1517001) on vitronectin (Thermo Fisher Scientific, A14700) pre-coated plates at 37 °C with 5% CO2. The Rho-associated protein kinase (ROCK) inhibitor Y-276325 (5 μM; Selleck Chemicals, S1049) was added to the E8 medium the first day after passaging or thawing of hESCs.

#### hESC-directed pancreatic differentiation

hESCs were seeded at a density of 2.3 × 10^5^ cells/cm^2^ on vitronectin-coated plates in E8 medium with 10 μM Y-27632. After 24 hours, cells were washed with PBS and differentiated to DE (stage 1), primitive gut tube (stage 2), PP1 (stage 3) and PP2 (stage 4) stages following previously described 4-stage protocol^40^. In brief, stage 1 (3 d): S1/2 medium supplemented with 100 ng ml^−1^ Activin A (Bon Opus Biosciences) and 5 μM CHIR99021 (04-0004-10, Stemgent) for 1 d. S1/2 medium supplemented with 100 ng ml^−1^Activin A for the next 2 d. Stage 2 (2 d): S1/2 medium supplemented with 50 ng ml^−1^ KGF (AF-100-19, PeproTech) and 0.25 mM vitamin C (VitC) (Sigma-Aldrich, A4544). Stage 3 (2 d): S3/4 medium supplemented with 50 ng ml^−1^ KGF, 0.25 mM VitC and 1 μM retinoic acid (R2625, MilliporeSigma). Stage 4 (4 d): S3/4 medium supplemented with 50 ng ml^−1^ KGF, 0.1 μM retinoic acid, 200 nM LDN (Stemgent, 04-0019), 0.25 μM SANT-1 (Sigma, S4572), 0.25 mM VitC and 200 nM TPB (EMD Millipore, 565740). The base differentiation medium formulations used in each stage were as follows. S1/2 medium: 500 ml MCDB 131 (15-100-CV, Cellgro) supplemented with 2 ml 45% glucose (G7528, MilliporeSigma), 0.75 g sodium bicarbonate (S5761, MilliporeSigma), 2.5 g BSA (68700, Proliant), 5 ml GlutaMAX (35050079, Invitrogen). S3/4 medium: 500 ml MCDB 131 supplemented with 0.52 ml 45% glucose, 0.875 g sodium bicarbonate, 10 g BSA, 2.5 ml ITS-X, 5 ml GlutaMAX.

#### Cell infection with LARRY barcode virus

Individual LARRY barcode constructs were cloned from the LARRY barcode library (Addgene:140024) and transfected to 293T cells to generate lentivirus. Next, each KO and WT hESC clone was infected with a unique LARRY barcode at low MOI. One week after lentiviral infection, the barcoded cells, which expressed GFP, were sorted out and cultured in E8 medium as described in previous section.

#### Pooled single-cell RNA-seq

One day before differentiation, each of 10 hESC barcoded clones were counted, mixed at the same cell number ratio, and then seeded at a density of 2.3 × 10^5^ cells/cm^2^ onto a 12-well cell culture plate. At DE and PP2 stages, pooled differentiating cells were dissociated into single cell suspension by TrypLE Select for 5 min at 37 °C. Cells were then stored in BAMBANKER™ freezing medium for future experiments. For scRNA-seq, frozen cells were thawed and sorted to collect live GFP+ cells. Cellular suspensions were then loaded on a Chromium Controller following the manufacturer’s instructions (10x Genomics Chromium Single Cell 3′ Reagent Kit v3.1 User Guide). cDNA libraries and targeted LARRY barcode libraries were generated separately using 10ul cDNA each. cDNA libraries were made under manufacturer’s instructions and targeted LARRY barcode libraries were amplified using specific primers (F: CTACACGACGCTCTTCCGATCT; R: GTGACTGGAGTTCAGACGTGTGCTCTTCCGATCTtaaccgttgctaggagagaccataT).

#### Data analysis

The sequencing data which included transcriptome and LARRY barcode libraries, underwent demultiplexing and processing via Cell Ranger (version 6.1.2). Subsequent analysis was conducted using the Seurat R package (version 4.3.1). Quality control measures were implemented to ensure robust data analysis. Cells were excluded if the number of expressed genes exceeded 7,000 or fell below 200. Additionally, cells were removed if the percentage of mitochondrial read exceeded 20%.

Singlet cells were defined by considering the highest feature barcode count, ensuring it was at least twice as large as the second highest feature barcode count. Single cells containing more than one detected barcode sequence were excluded from the dataset. This process resulted in a final set of 20,678 cells for downstream analysis. After quality control measures, the count matrix underwent normalization and scaling. PCA was performed using the top 2,000 highly variable genes. Subsequently, clustering and UMAP embeddings were generated using default parameters to elucidate the underlying structure and relationships within the dataset.

#### Flow cytometry

Cells were dissociated using TrypLE Select and resuspended in FACS buffer (5% FBS in PBS). Live/Dead Fixable Violet cell stain (Invitrogen, L34955) was used to discriminate dead cells from live cells. Permeabilization/fixation was performed at room temperature for 1 h. Antibody staining was performed in permeabilization buffer. Antibodies for this study include HNF4A, Novus Biologicals, NBP2-67679, 1:200; PDX1, R&D Systems, AF2419, 1:500, Donkey anti-Rabbit IgG (H+L) Highly Cross-Adsorbed Secondary Antibody, Thermo Fisher Scientific, 1:500; Donkey anti-goat IgG (H+L) Highly Cross-Adsorbed Secondary Antibody, Thermo Fisher Scientific, 1:500. Cells were then analysed using BD LSRFortessa. Flow cytometry analysis and figures were generated using FlowJo v.10.

## Acknowledgements

The authors thank all members from the Li and Huangfu laboratory for comments and discussions. The authors thank Jake P. Taylor-King for discussions. This study is supported by NIH R01 HG010753, HL168174 (to W.L., B.S., L.C.), District of Columbia Center for AIDS (DC-CFAR) Research Transitioning Investigator Award (AI117970, to W.L.), and startup support from the Center for Genetic Medicine Research at Children’s National Hospital. D.H is supported by NIH UM1 HG012654, U01 HG012051. J.J. Li is supported by National Science Foundation DBI-1846216 and DMS-2113754, NIH R35 GM140888, Johnson and Johnson WiSTEM2D Award, Sloan Research Fellowship, UCLA David Geffen School of Medicine W.M. Keck Foundation Junior Faculty Award, and Chan-Zuckerberg Initiative Single-Cell Biology Data Insights [Silicon Valley Community Foundation Grant Number: 2022-249355]. W.D. and R.F.S. is supported by the Howard Hughes Medical Institute.

## Author contributions

W.L. conceived the project. W.L. and B.S. developed the method. W.L., B.S., and D.L. designed and performed the experiments and analyzed the data. W.D., N.M. and B.S. performed and analyzed HIV Perturb-seq under the supervision of J.M.S., R.S. and W.L. B.S., Q.W. and D.S. performed synthetic experiments under the supervision of W.L. and J.J.L. D.L., D.Y., B.W. and B.R. generated pancreatic differentiation dataset and performed validations under the supervision of D.H. A.K., A.V., N.U., and A.L. generated and analyzed genome-scale Perturb-seq under the supervision of T.B. X.C., L.C. and Y.D. performed the analysis and interpretation of the results. W.L., and B.S. wrote the manuscript with input from all the authors. W.L. T.B., J.J.L., R.S. and D.H. supervised the study.

## Data and materials availability

The Perturb-seq scRNA-seq data have been deposited to Gene Expression Omnibus (GEO) under the accession number GSE247601. The source code of the PS method, and the documentation and demos are available on GitHub: (https://github.com/davidliwei/PS).

## Supplementary Figure Legends

**Supplementary Figure S1. Benchmark different methods using simulated and real datasets. a**, Steps to generate simulated datasets using scDesign3 from a real scRNA-seq dataset that knocks out Nelfb gene. **b-e**, The score distribution of scMAGeCK-PS and mixscape using different DEGs and different values of true efficiencies. **f-g**, Similar with Figure 1f-g, but using a published high MOI Perturb-seq dataset in the same study. **h-i**, Benchmark results of different methods on another published CRISPRi-based Perturb-seq, where mismatches are introduced into guides to attenuate perturbation effects. The Pearson correlation coefficients (PCCs) between the predicted scores of each method and the expressions of perturbed genes are reported for every perturbed gene (**h**), and between the predicted scores and predicted sgRNA activities (**i**), using the prediction methods provided in the original study^21^.

**Supplementary Figure S2. A genome-scale Perturb-seq. a-b**, The distribution of scMAGeCK-PS and mixscape predicted scores of top hits including CD247 (**a**) and LCK (**b**) in the pooled screen. **c**, The correlation between PSs and perturbed gene expression.

**Supplementary Figure S3. Predictions of PDL1 protein expression from a published ECCITE-seq dataset**. The ROC curve, the correlations between scMAGeCK-PS results with PDL1 protein expression, and the correlations between mixscape results with PDL1 protein expression are reported for each gene. The correlations are separated by classifications of each single cell: NP (non-perturbed), defined as mixscape score <=0.5; and KO (knockout), defined as mixscape score >0.5. For a fair comparison, we used mixscape classification results to plot PSs (mid panel).

**Supplementary Figure S4. Buffered genes and sensitive genes. a**, RPL4, a buffered gene. **b-c**, HSPA5 and GATA1, two sensitive genes. **d**, A gene (BRD4) whose expression has no correlation with PS.

**Supplementary Figure S5. The log fold changes of gene expressions upon perturbing genes within the same protein complex**, including ribosomal subunits (**a**), RNA polymerase (**b**) and mediator complex (**c**) in essential gene Perturb-seq. **(d)** The log fold changes of proteosome gene expressions (columns) upon perturbing proteasome genes (rows) from the genome-scale Perturb-seq.

**Supplementary Figure S6. HIV Perturb-seq. a**, The number of genes (nFeature_RNA), UMI counts (nCount_RNA) and the fraction of mitochondrial RNAs in three different conditions. **b**, Clustering results. **c**, Enriched Gene Ontology (GO) terms of cluster 8. **d**, The distribution of BRD4-targeting gRNAs. **e**, The expression distribution of BRD4 signature genes in cluster 8 vs other clusters. Only cells express BRD4-targeting gRNAs are included. **f**, Differential expression results between BRD4 PS+ cells vs BRD4 PS-cells.

**Supplementary Figure S7. HIV Perturb-seq. a**, The expressions of CCNT1 (left) and CCNT1-targeting gRNAs (right). **b**, Differential expression results between CCNT1-targeting cells and non-targeting control cells in two different cell states. **c**, The expressions of NFKB1. **d**, The quantitative perturbation-expression relationship between GFP and CCNT1 PS, similar with Figure 4e.

**Supplementary Figure S8. Cell type assignment based on known expression markers of different cell types in pancreatic differentiation scRNA-seq**.

**Supplementary Figure S9. DEG analysis**. a**-b**, The distribution of FOXA1 PSs across two different clones. **c**, The expression pattern of FOXA1. **d-e**, The DEG analysis results of CCDC6 knockout clones vs. wild-type clones in different cell types. **f**, The overlap of statistically significant DEGs in DE and LV/DUO cell types.

**Supplementary Figure S10. Different CCDC6 functions. a-b**, The two patterns of CCDC6 PSs in LV/DUO (**a**) and DE in transition (**b**) cell types. **c-f**, Additional enriched terms using Enrichr on DEGs of CCDC6 knockout.

**Supplementary Figure S11. Flow cytometry analysis of PDX1 and HNF4A expression upon CCDC6 knockout**. One representative plots of three biological replicates are shown.

**Supplementary Table S1. Genome-scale Perturb-seq library design. Supplementary Table S2. HIV Perturb-seq library design**.

**Supplementary Table S3. Sequencing summary of HIV Perturb-seq**.

**Supplementary Table S4. Genotype summary of 10-clone scRNA-seq pancreatic differentiation dataset**.

## References

1 High-content CRISPR screening. Nat Rev Methods Primers 2022;2:. 10.1038/s43586-022-00098-7.

2 Srivatsan SR, McFaline-Figueroa JL, Ramani V, Saunders L, Cao J, Packer J, et al. Massively multiplex chemical transcriptomics at single-cell resolution. Science 2020;367:45–51.

3 Dixit A, Parnas O, Li B, Chen J, Fulco CP, Jerby-Arnon L, et al. Perturb-Seq: Dissecting Molecular Circuits with Scalable Single-Cell RNA Profiling of Pooled Genetic Screens. Cell 2016;167:1853–1866.e17.

4 Adamson B, Norman TM, Jost M, Cho MY, Nuñez JK, Chen Y, et al. A Multiplexed Single-Cell CRISPR Screening Platform Enables Systematic Dissection of the Unfolded Protein Response. Cell 2016;167:1867–1882.e21.

5 Jaitin DA, Weiner A, Yofe I, Lara-Astiaso D, Keren-Shaul H, David E, et al. Dissecting Immune Circuits by Linking CRISPR-Pooled Screens with Single-Cell RNA-Seq. Cell 2016;167:1883–1896.e15.

6 Datlinger P, Rendeiro AF, Schmidl C, Krausgruber T, Traxler P, Klughammer J, et al. Pooled CRISPR screening with single-cell transcriptome readout. Nat Methods 2017;14:297–301.

7 Xie S, Duan J, Li B, Zhou P, Hon GC. Multiplexed Engineering and Analysis of Combinatorial Enhancer Activity in Single Cells. Mol Cell 2017;66:285–299.e5.

8 Rubin AJ, Parker KR, Satpathy AT, Qi Y, Wu B, Ong AJ, et al. Coupled Single-Cell CRISPR Screening and Epigenomic Profiling Reveals Causal Gene Regulatory Networks. Cell 2019;176:361–376.e17.

9 Liscovitch-Brauer N, Montalbano A, Deng J, Méndez-Mancilla A, Wessels H-H, Moss NG, et al. Profiling the genetic determinants of chromatin accessibility with scalable single-cell CRISPR screens. Nat Biotechnol 2021;39:1270–7.

10 Dhainaut M, Rose SA, Akturk G, Wroblewska A, Nielsen SR, Park ES, et al. Spatial CRISPR genomics identifies regulators of the tumor microenvironment. Cell 2022;185:1223–1239.e20.

11 Norman TM, Horlbeck MA, Replogle JM, Ge AY, Xu A, Jost M, et al. Exploring genetic interaction manifolds constructed from rich single-cell phenotypes. Science 2019;365:786–93.

12 Replogle JM, Norman TM, Xu A, Hussmann JA, Chen J, Cogan JZ, et al. Combinatorial single-cell CRISPR screens by direct guide RNA capture and targeted sequencing. Nat Biotechnol 2020;38:954–61.

13 Wessels H-H, Méndez-Mancilla A, Hao Y, Papalexi E, Mauck WM 3rd, Lu L, et al. Efficient combinatorial targeting of RNA transcripts in single cells with Cas13 RNA Perturb-seq. Nat Methods 2023;20:86–94.

14 Hill AJ, McFaline-Figueroa JL, Starita LM, Gasperini MJ, Matreyek KA, Packer J, et al. On the design of CRISPR-based single-cell molecular screens. Nat Methods 2018;15:271–4.

15 Yang L, Zhu Y, Yu H, Cheng X, Chen S, Chu Y, et al. scMAGeCK links genotypes with multiple phenotypes in single-cell CRISPR screens. Genome Biol 2020;21:19.

16 Papalexi E, Mimitou EP, Butler AW, Foster S, Bracken B, Mauck WM 3rd, et al. Characterizing the molecular regulation of inhibitory immune checkpoints with multimodal single-cell screens. Nat Genet 2021;53:322–31.

17 Tsuchida CA, Brandes N, Bueno R, Trinidad M, Mazumder T, Yu B, et al. Mitigation of chromosome loss in clinical CRISPR-Cas9-engineered T cells. Cell 2023;186:4567–4582.e20.

18 Song D, Wang Q, Yan G, Liu T, Sun T, Li JJ. scDesign3 generates realistic in silico data for multimodal single-cell and spatial omics. Nat Biotechnol 2023. 10.1038/s41587-023-01772-1.

19 Wu B, Zhang X, Chiang H-C, Pan H, Yuan B, Mitra P, et al. RNA polymerase II pausing factor NELF in CD8+ T cells promotes antitumor immunity. Nat Commun 2022;13:2155.

20 Gasperini M, Hill AJ, McFaline-Figueroa JL, Martin B, Kim S, Zhang MD, et al. A Genome-wide Framework for Mapping Gene Regulation via Cellular Genetic Screens. Cell 2019;176:1516.

21 Jost M, Santos DA, Saunders RA, Horlbeck MA, Hawkins JS, Scaria SM, et al. Titrating gene expression using libraries of systematically attenuated CRISPR guide RNAs. Nat Biotechnol 2020;38:355–64.

22 Schraivogel D, Gschwind AR, Milbank JH, Leonce DR, Jakob P, Mathur L, et al. Targeted Perturb-seq enables genome-scale genetic screens in single cells. Nat Methods 2020;17:629–35.

23 Shifrut E, Carnevale J, Tobin V, Roth TL, Woo JM, Bui CT, et al. Genome-wide CRISPR screens in primary human T cells reveal key regulators of immune function. Cell 2018;175:1958–1971.e15.

24 Wessels H-H, Stirn A, Méndez-Mancilla A, Kim EJ, Hart SK, Knowles DA, et al. Prediction of ontarget and off-target activity of CRISPR-Cas13d guide RNAs using deep learning. Nat Biotechnol 2023. 10.1038/s41587-023-01830-8.

25 Zhang J, Bu X, Wang H, Zhu Y, Geng Y, Nihira NT, et al. Cyclin D–CDK4 kinase destabilizes PD-L1 via cullin 3–SPOP to control cancer immune surveillance. Nature 2018;553:91–5.

26 Naqvi S, Kim S, Hoskens H, Matthews HS, Spritz RA, Klein OD, et al. Precise modulation of transcription factor levels identifies features underlying dosage sensitivity. Nat Genet 2023;55:841–51.

27 Replogle JM, Saunders RA, Pogson AN, Hussmann JA, Lenail A, Guna A, et al. Mapping information-rich genotype-phenotype landscapes with genome-scale Perturb-seq. Cell 2022;185:2559–2575.e28.

28 Radhakrishnan SK, Lee CS, Young P, Beskow A, Chan JY, Deshaies RJ. Transcription factor Nrf1 mediates the proteasome recovery pathway after proteasome inhibition in mammalian cells. Mol Cell 2010;38:17–28.

29 Dai W, Wu F, McMyn N, Song B, Walker-Sperling VE, Varriale J, et al. Genome-wide CRISPR screens identify combinations of candidate latency reversing agents for targeting the latent HIV-1 reservoir. Sci Transl Med 2022;14:eabh3351.

30 Yin M, Guo Y, Hu R, Cai WL, Li Y, Pei S, et al. Potent BRD4 inhibitor suppresses cancer cell-macrophage interaction. Nat Commun 2020;11:1833.

31 Tan Y-F, Wang M, Chen Z-Y, Wang L, Liu X-H. Inhibition of BRD4 prevents proliferation and epithelial-mesenchymal transition in renal cell carcinoma via NLRP3 inflammasome-induced pyroptosis. Cell Death Dis 2020;11:239.

32 Shu S, Wu H-J, Ge JY, Zeid R, Harris IS, Jovanović B, et al. Synthetic Lethal and Resistance Interactions with BET Bromodomain Inhibitors in Triple-Negative Breast Cancer. Mol Cell 2020;78:1096–1113.e8.

33 Li Z, Guo J, Wu Y, Zhou Q. The BET bromodomain inhibitor JQ1 activates HIV latency through antagonizing Brd4 inhibition of Tat-transactivation. Nucleic Acids Res 2013;41:277–87.

34 Mbonye U, Kizito F, Karn J. New insights into transcription elongation control of HIV-1 latency and rebound. Trends Immunol 2023;44:60–71.

35 Wei P, Garber ME, Fang S-M, Fischer WH, Jones KA. A novel CDK9-associated C-type cyclin interacts directly with HIV-1 tat and mediates its high-affinity, loop-specific binding to TAR RNA. Cell 1998;92:451–62.

36 Peng J, Zhu Y, Milton JT, Price DH. Identification of multiple cyclin subunits of human P-TEFb. Genes Dev 1998;12:755–62.

37 Stoeckius M, Zheng S, Houck-Loomis B, Hao S, Yeung BZ, Mauck WM 3rd, et al. Cell Hashing with barcoded antibodies enables multiplexing and doublet detection for single cell genomics. Genome Biol 2018;19:224.

38 Weinreb C, Rodriguez-Fraticelli A, Camargo FD, Klein AM. Lineage tracing on transcriptional landscapes links state to fate during differentiation. Science 2020;367:eaaw3381.

39 Vitak SA, Torkenczy KA, Rosenkrantz JL, Fields AJ, Christiansen L, Wong MH, et al. Sequencing thousands of single-cell genomes with combinatorial indexing. Nat Methods 2017;14:302–8.

40 Yang D, Cho H, Tayyebi Z, Shukla A, Luo R, Dixon G, et al. CRISPR screening uncovers a central requirement for HHEX in pancreatic lineage commitment and plasticity restriction. Nat Cell Biol 2022;24:1064–76.

41 Rosen BP, Li QV, Cho H, Liu D, Yang D, Graff S, et al. Parallel genome-scale CRISPR screens distinguish pluripotency and self-renewal. BioRxivorg 2023. 10.1101/2023.05.03.539283.

42 Li QV, Dixon G, Verma N, Rosen BP, Gordillo M, Luo R, et al. Genome-scale screens identify JNK-JUN signaling as a barrier for pluripotency exit and endoderm differentiation. Nat Genet 2019;51:999–1010.

43 Thanasopoulou A, Stravopodis DJ, Dimas KS, Schwaller J, Anastasiadou E. Loss of CCDC6 affects cell cycle through impaired intra-S-phase checkpoint control. PLoS One 2012;7:e31007.

44 Morra F, Luise C, Merolla F, Poser I, Visconti R, Ilardi G, et al. FBXW7 and USP7 regulate CCDC6 turnover during the cell cycle and affect cancer drugs susceptibility in NSCLC. Oncotarget 2015;6:12697–709.

45 Zhou Y, Luo K, Liang L, Chen M, He X. A new Bayesian factor analysis method improves detection of genes and biological processes affected by perturbations in single-cell CRISPR screening. Nat Methods 2023. 10.1038/s41592-023-02017-4.

46 Dong M, Wang B, Wei J, de O Fonseca AH, Perry CJ, Frey A, et al. Causal identification of singlecell experimental perturbation effects with CINEMA-OT. Nat Methods 2023;20:1769–79.

47 Hart T, Brown KR, Sircoulomb F, Rottapel R, Moffat J. Measuring error rates in genomic perturbation screens: gold standards for human functional genomics. Mol Syst Biol 2014;10:733.

48 Morgens DW, Deans RM, Li A, Bassik MC. Systematic comparison of CRISPR/Cas9 and RNAi screens for essential genes. Nat Biotechnol 2016;34:634–6.

49 Bunne C, Stark SG, Gut G, Del Castillo JS, Levesque M, Lehmann K-V, et al. Learning single-cell perturbation responses using neural optimal transport. Nat Methods 2023;20:1759–68.

50 Li WV, Li JJ. An accurate and robust imputation method scImpute for single-cell RNA-seq data. Nat Commun 2018;9:997.

51 Lun ATL, Bach K, Marioni JC. Pooling across cells to normalize single-cell RNA sequencing data with many zero counts. Genome Biol 2016;17:75.

52 Sun T, Song D, Li WV, Li JJ. scDesign2: a transparent simulator that generates high-fidelity single-cell gene expression count data with gene correlations captured. Genome Biol 2021;22:163.

53 Hao Y, Hao S, Andersen-Nissen E, Mauck WM 3rd, Zheng S, Butler A, et al. Integrated analysis of multimodal single-cell data. Cell 2021;184:3573–3587.e29.

54 Lee K, Cho H, Rickert RW, Li QV, Pulecio J, Leslie CS, et al. FOXA2 is required for enhancer priming during pancreatic differentiation. Cell Rep 2019;28:382–393.e7.

