## Supplementary Figures for "Decoding Heterogenous Single-cell Perturbation Responses"

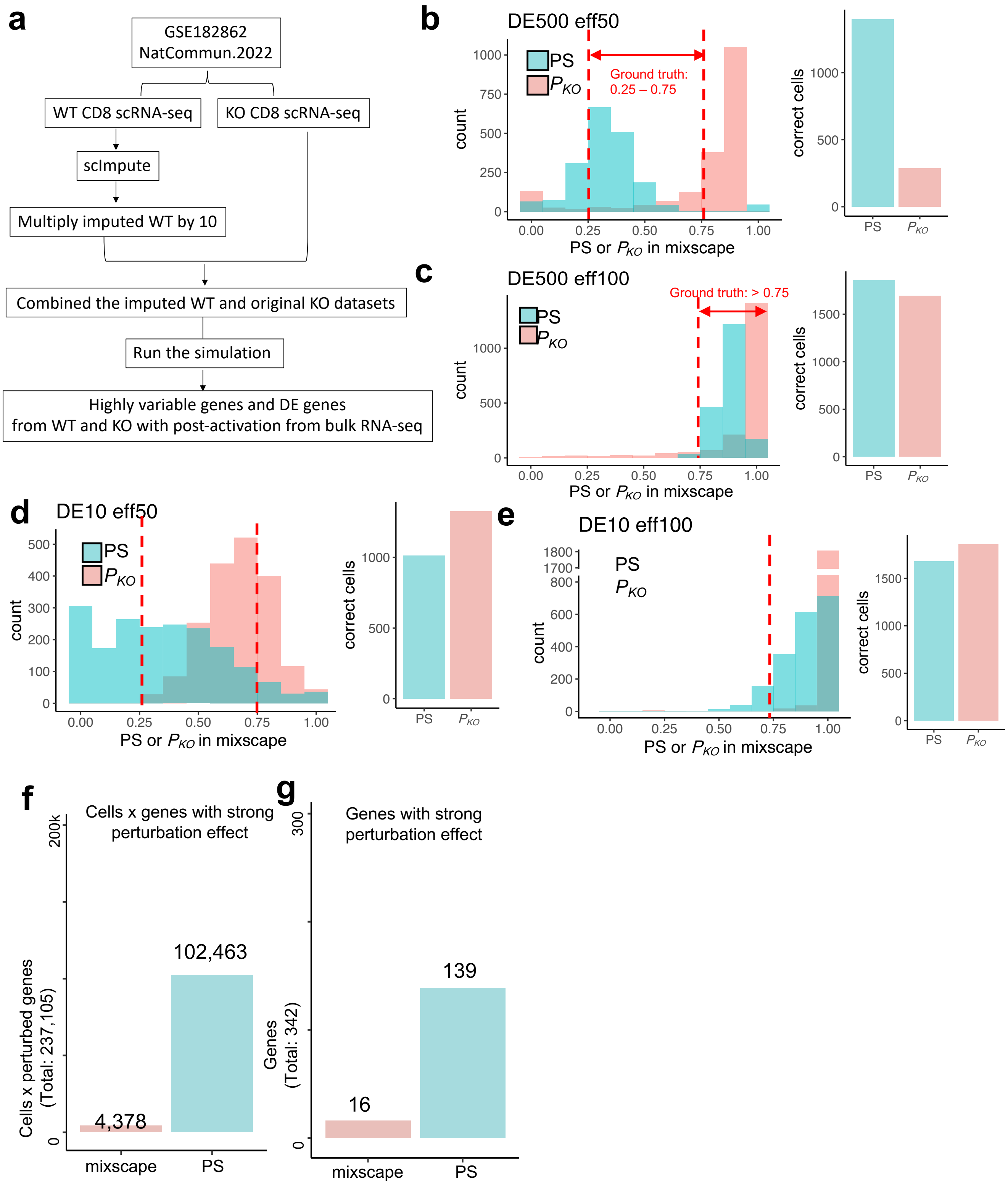
a
GSE182862
NatCommun.2022
WT CD8 scRNA-seq
KO CD8 scRNA-seq
scImpute
Multiply imputed WT by 10
Combined the imputed WT and original KO datasets
Run the simulation
Highly variable genes and DE genes
from WT and KO with post-activation from bulk RNA-seq
b
DE500 eff50
PS
PKO
Ground truth:
0.25 – 0.75
count
PS or PKO in mixscape
correct cells
c
DE500 eff100
Ground truth: > 0.75
d
DE10 eff50
e
DE10 eff100
f
Cells x genes with strong
perturbation effect
Cells x perturbed genes
(Total: 237,105)
102,463
4,378
mixscape
PS
g
Genes with strong
perturbation effect
Genes
(Total: 342)
139
16

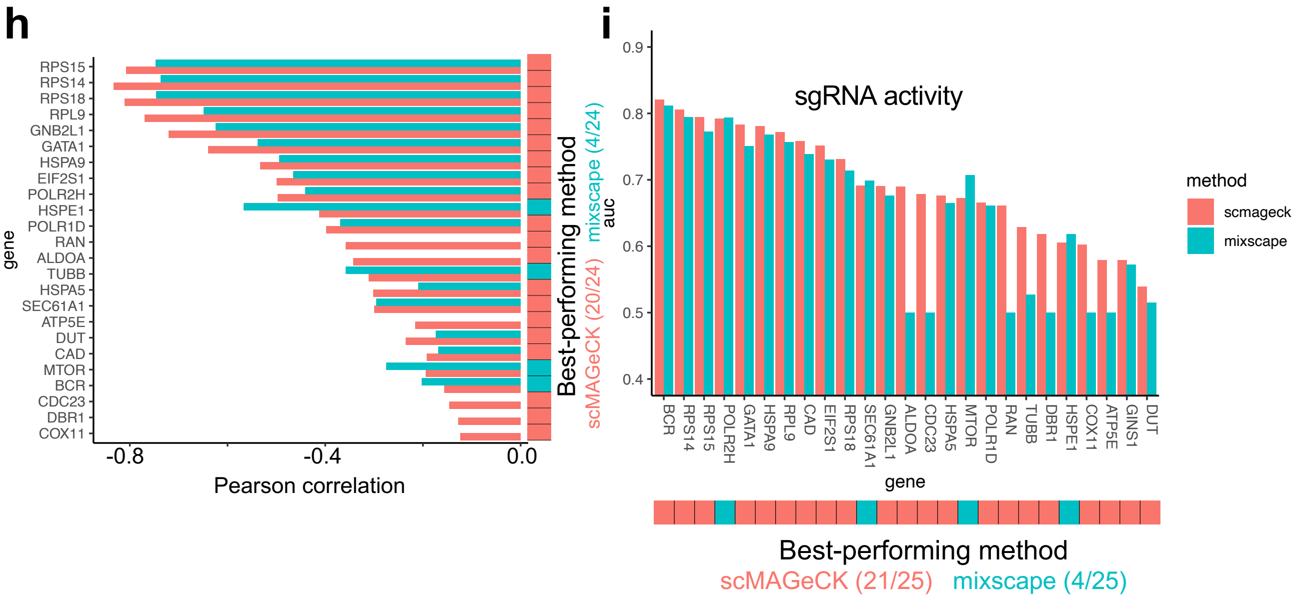


**Figure S1. Benchmarking.**

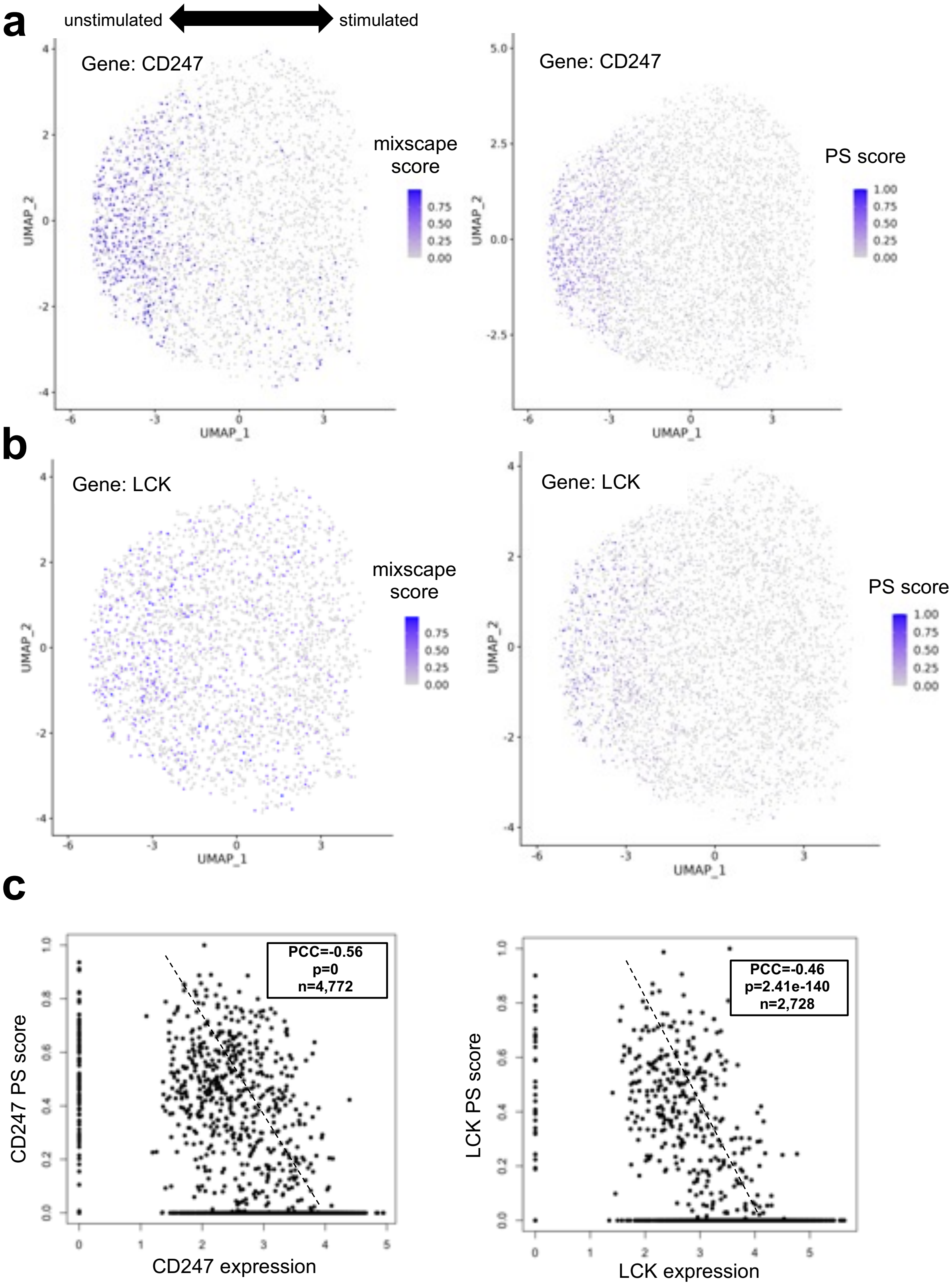


**Figure S2:** Genome-scale Perturb-seq

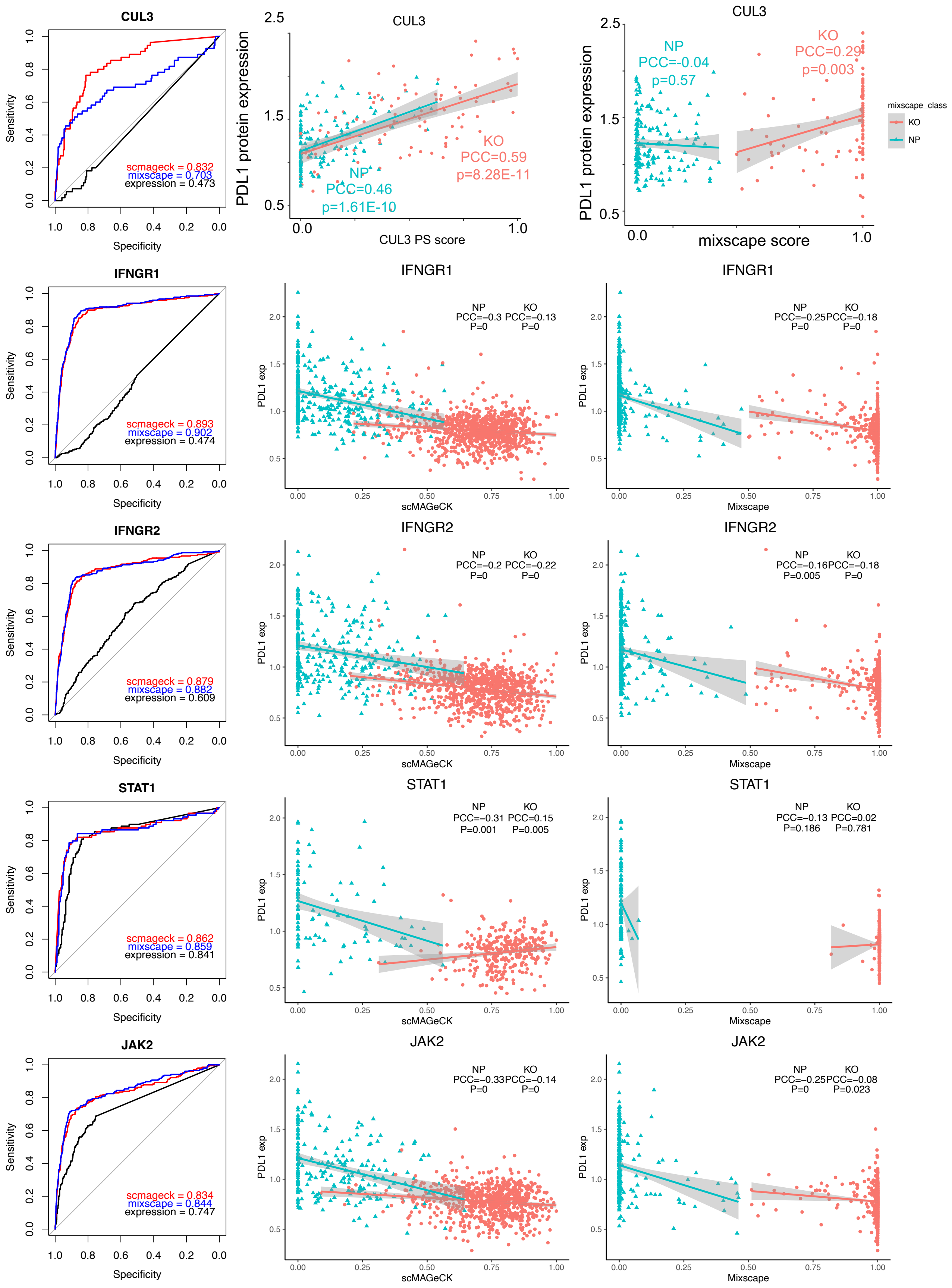
CUL3
Sensitivity
Specificity
scmageck = 0.832
mixscape = 0.703
expression = 0.473
CUL3
PDL1 protein expression
NP
PCC=0.46
p=1.61E-10
KO
PCC=0.59
p=8.28E-11
CUL3 PS score
CUL3
NP
PCC=-0.04
p=0.57
KO
PCC=0.29
p=0.003
mixscape score
mixscape_class
KO
NP
IFNGR1
scmageck = 0.893
mixscape = 0.902
expression = 0.474
IFNGR1
NP PCC=-0.3 P=0
KO PCC=-0.13 P=0
PDL1 exp
scMAGeCK
IFNGR1
NP PCC=-0.25 P=0
KO PCC=-0.18 P=0
Mixscape
IFNGR2
scmageck = 0.879
mixscape = 0.882
expression = 0.609
IFNGR2
NP PCC=-0.2 P=0
KO PCC=-0.22 P=0
IFNGR2
NP PCC=-0.16 P=0.005
KO PCC=-0.18 P=0
STAT1
scmageck = 0.862
mixscape = 0.859
expression = 0.841
STAT1
NP PCC=-0.31 P=0.001
KO PCC=0.15 P=0.005
STAT1
NP PCC=-0.13 P=0.186
KO PCC=0.02 P=0.781
JAK2
scmageck = 0.834
mixscape = 0.844
expression = 0.747
JAK2
NP PCC=-0.33 P=0
KO PCC=-0.14 P=0
JAK2
NP PCC=-0.25 P=0
KO PCC=-0.08 P=0.023

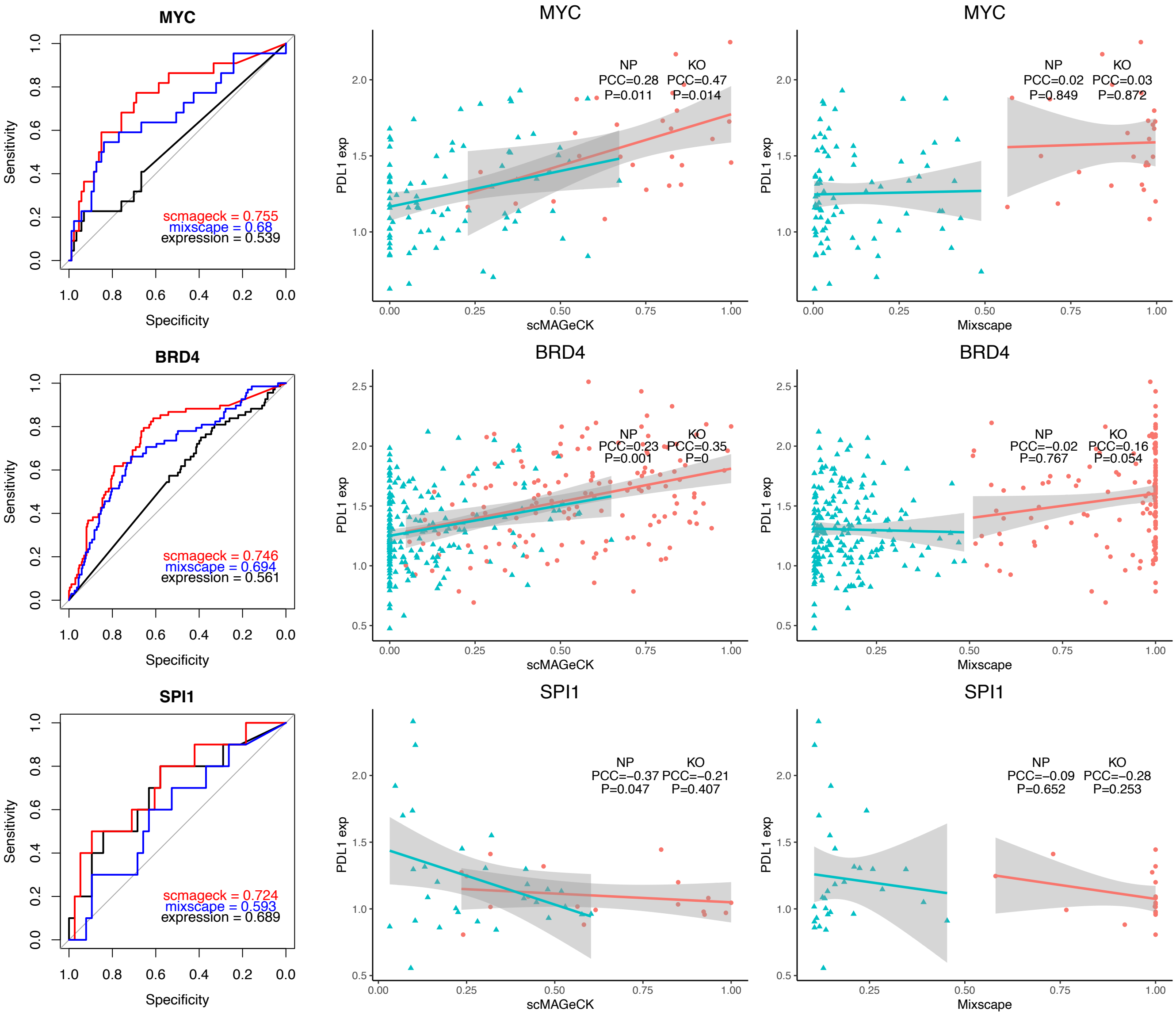


**Figure S3:** ECCITE-seq

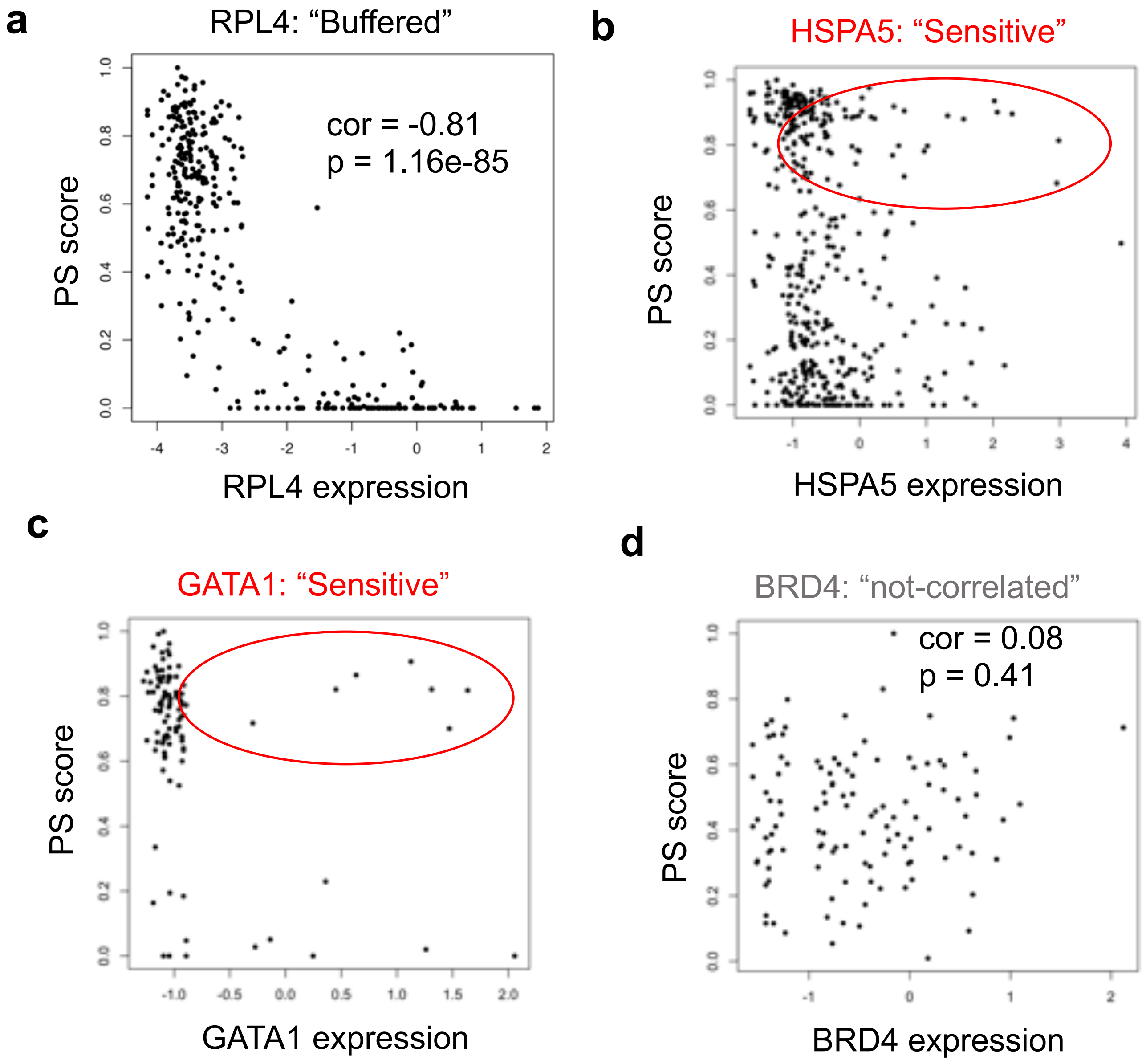


**Figure S4:** Buffered and sensitive gene

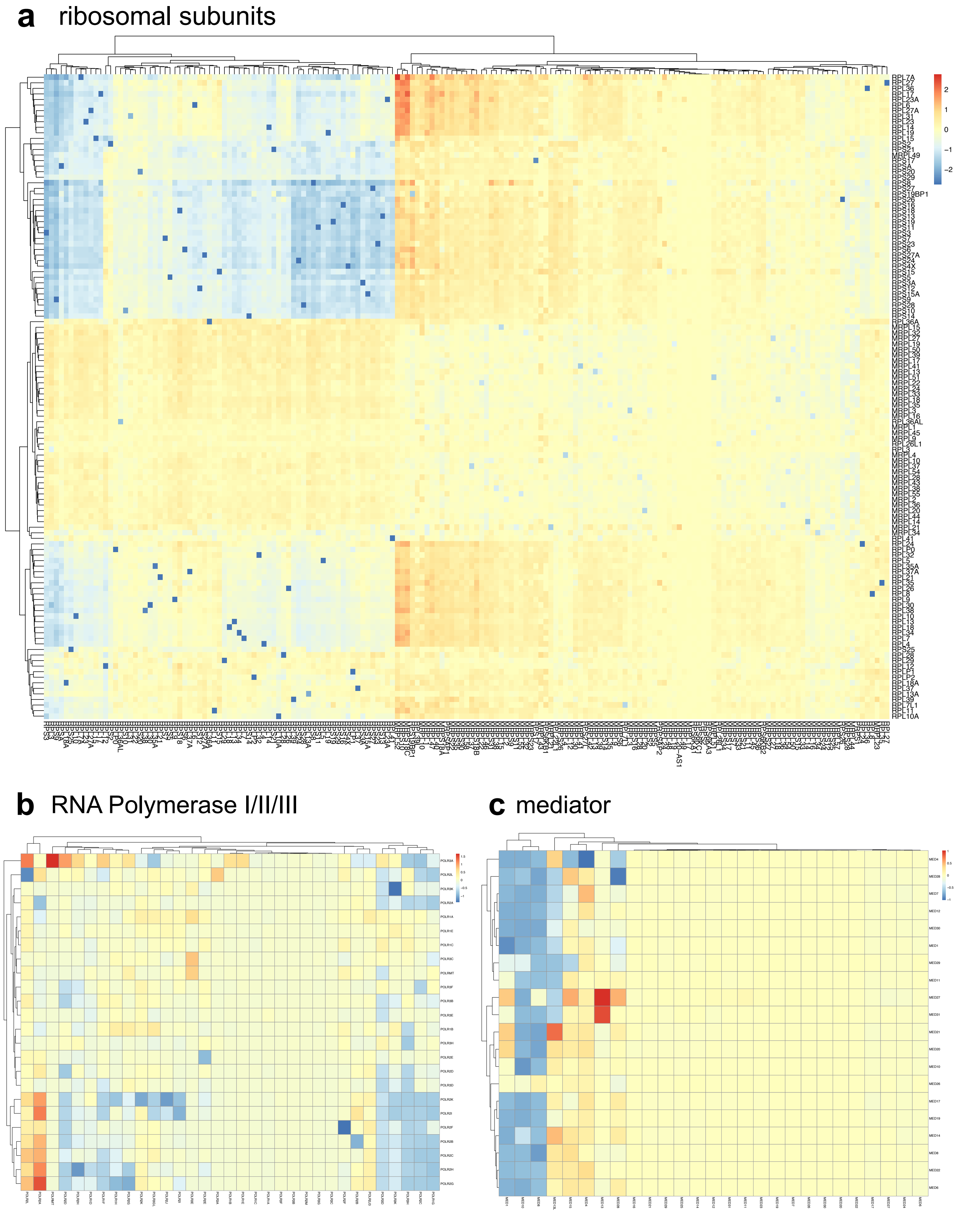
a ribosomal subunits
b RNA Polymerase I/II/III
c mediator

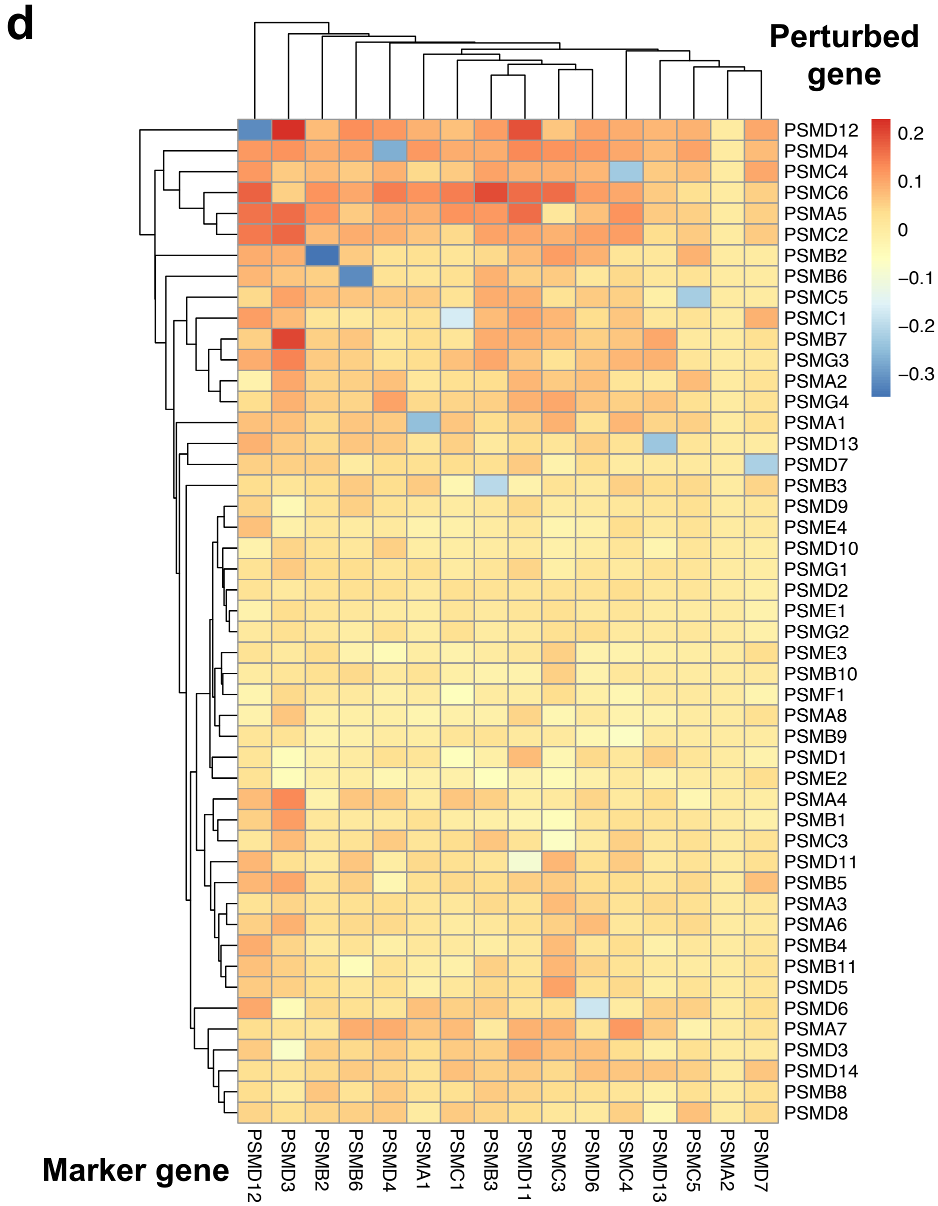


**Figure S5:** Expression compensation.

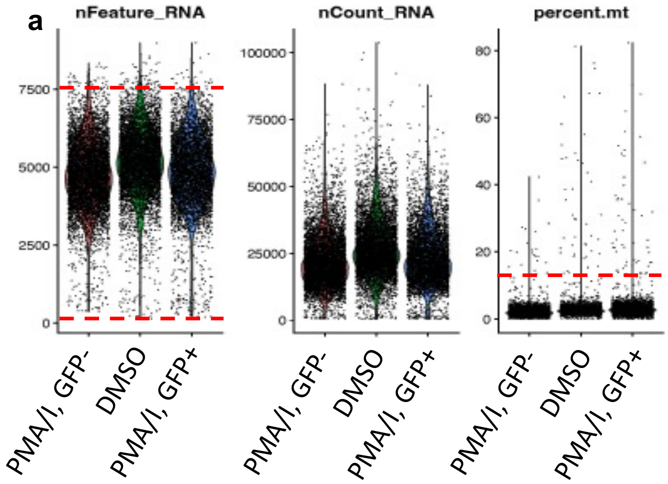


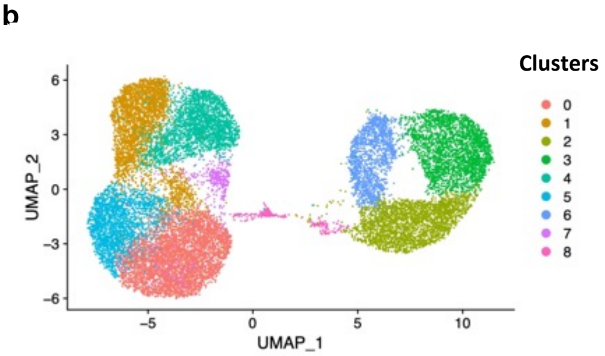


c

| Term | P-value | Adjusted P-value | Odds Ratio |
|---|---|---|---|
| Apoptosis | 2.16E-04 | 3.99E-03 | 6.26 |
| Hypoxia | 1.36E-04 | 3.99E-03 | 5.77 |
| Mitotic Spindle | 3.94E-03 | 3.73E-02 | 4.24 |
| TNF-alpha Signaling via NF-kB | 4.04E-03 | 3.73E-02 | 4.22 |

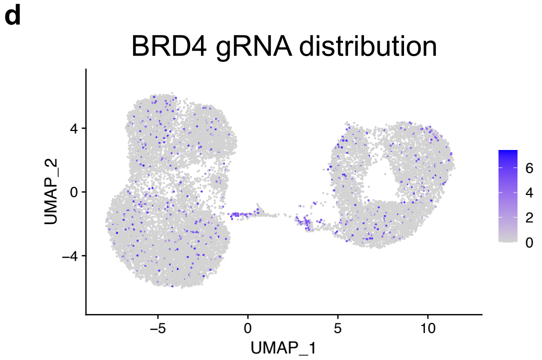


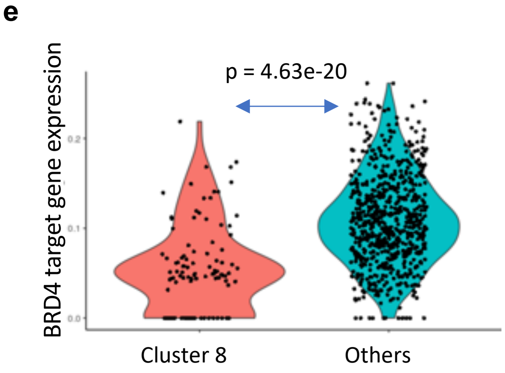


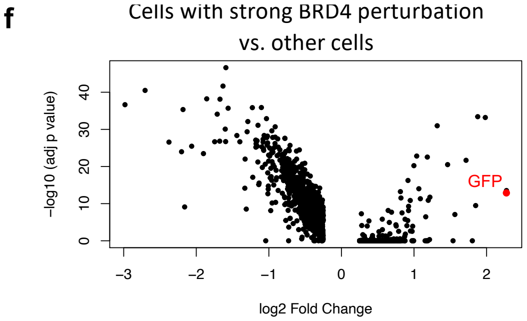


**Figure S6:** HIV Perturb-seq

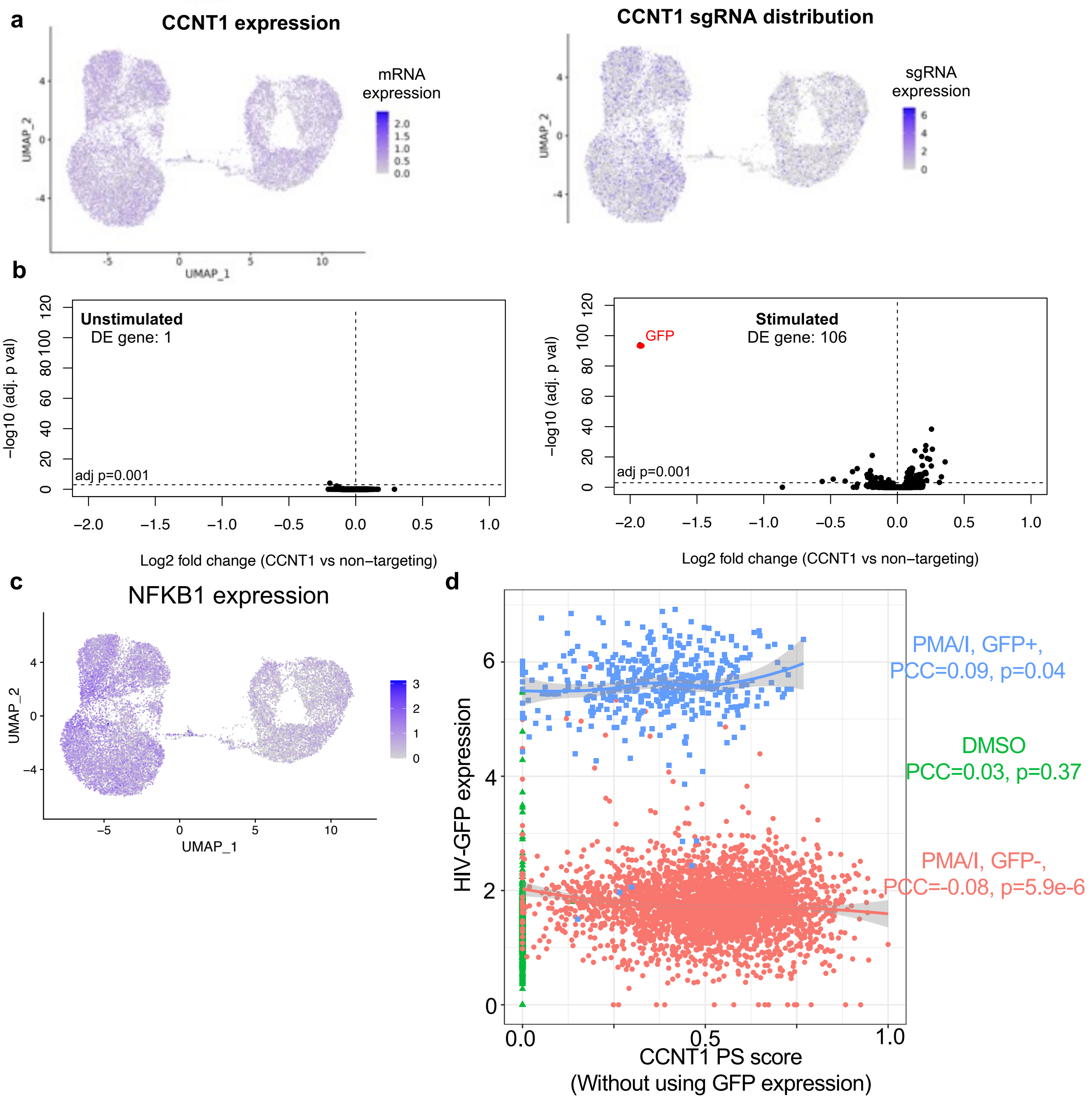


**Figure S7:** HIV Perturb-seq

**DE in transition: POU5F1, NANOG, SOX2** Cluster 2

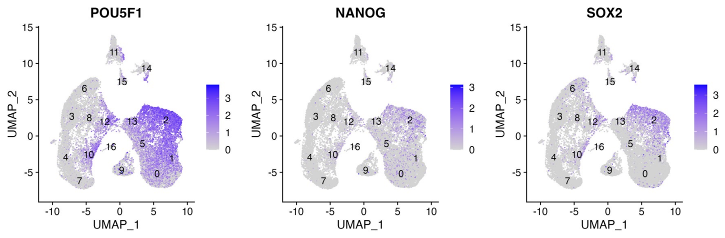


**Definitive endoderm: GATA6, SOX17, FOXA2, CXCR4** Clusters 0, 1, 5, 13

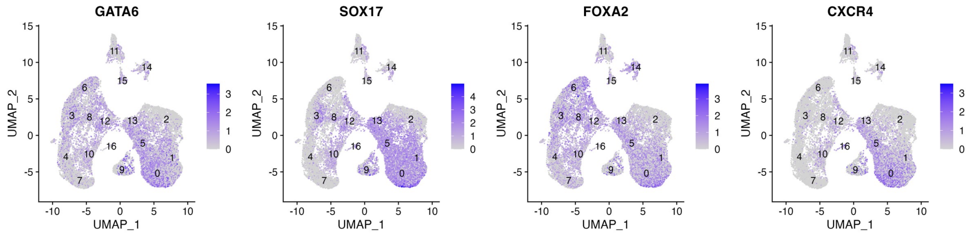


**PP1: Early-stage Pancreatic progenitor (NKX6.1-): PDX1, SOX9, FOXA2, HHEX**

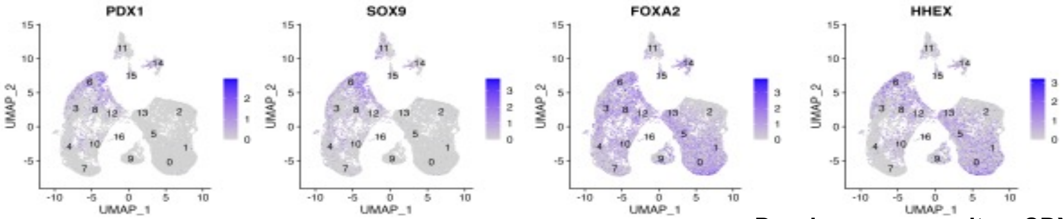


**PP2: Late-stage Pancreatic progenitor (NKX6.1+): PDX1, NKX6.1** Cluster 6

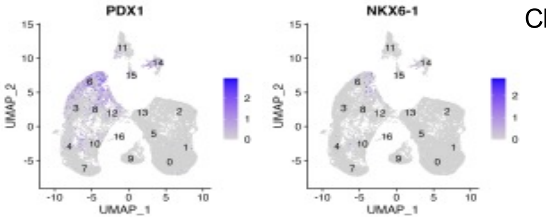


**Duodenum progenitor: CDX2, LGALS3**

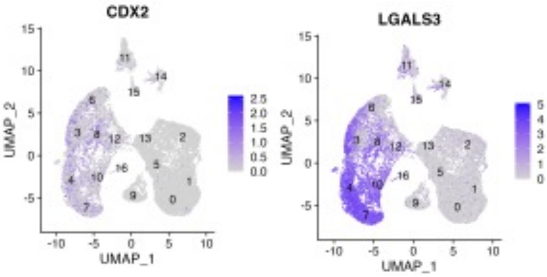


**Endocrine progenitor: NEUROG3, CHGA, NKX2.2** Cluster 14

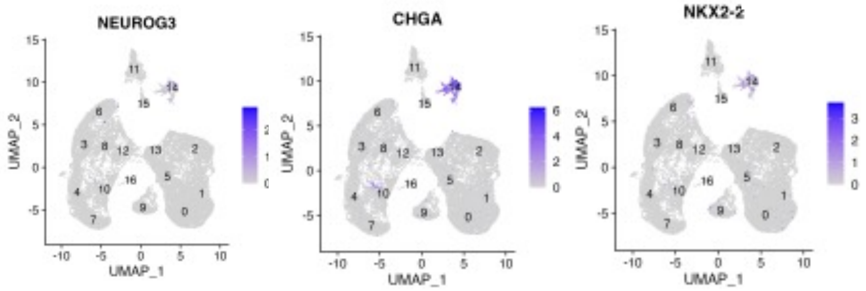


**Liver progenitor: AFP, APOA2, FABP1** Clusters 4, 7, 10

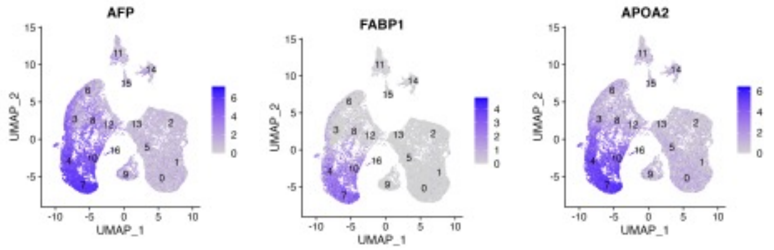


**Figure S8:** 10-clone scRNA-seq

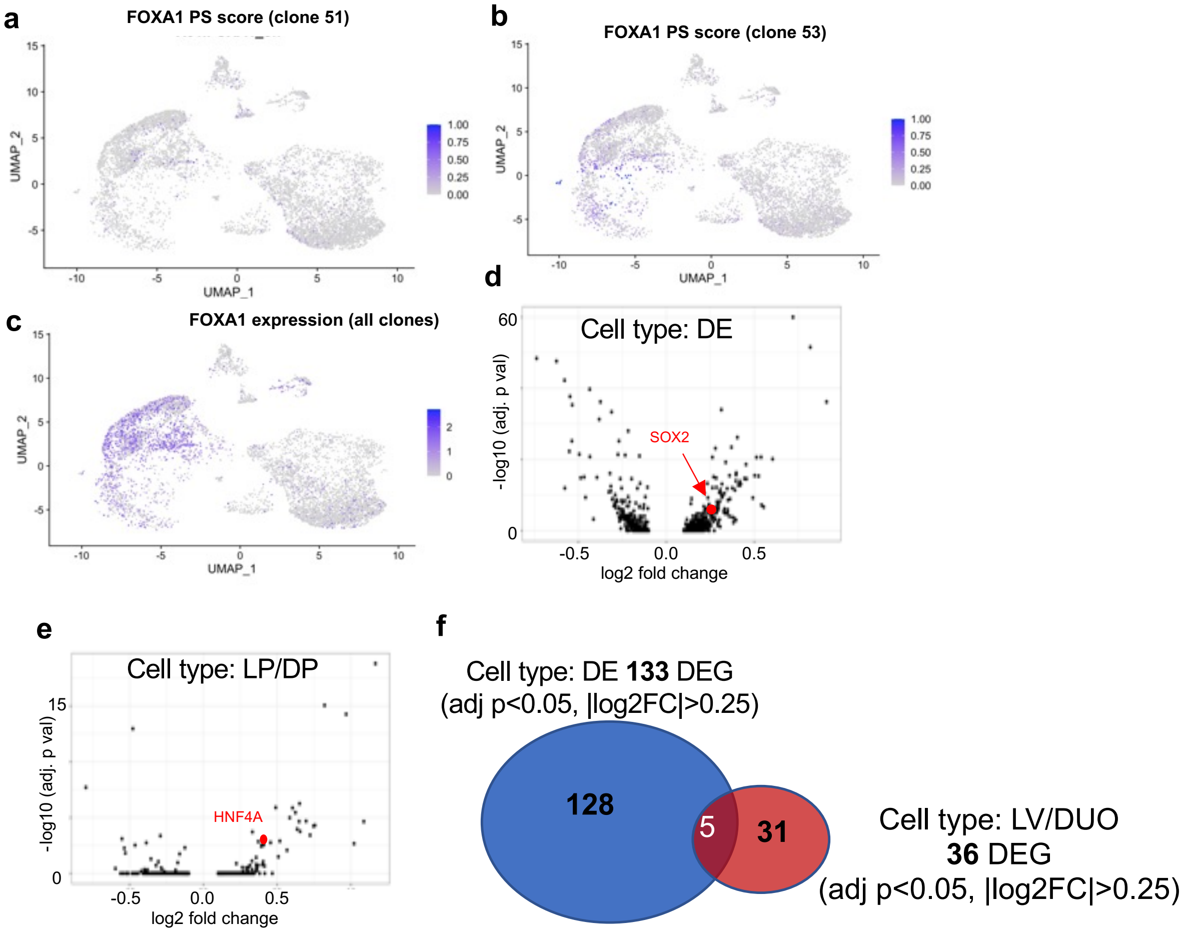


**Figure S9:** 10-clone scRNA-seq

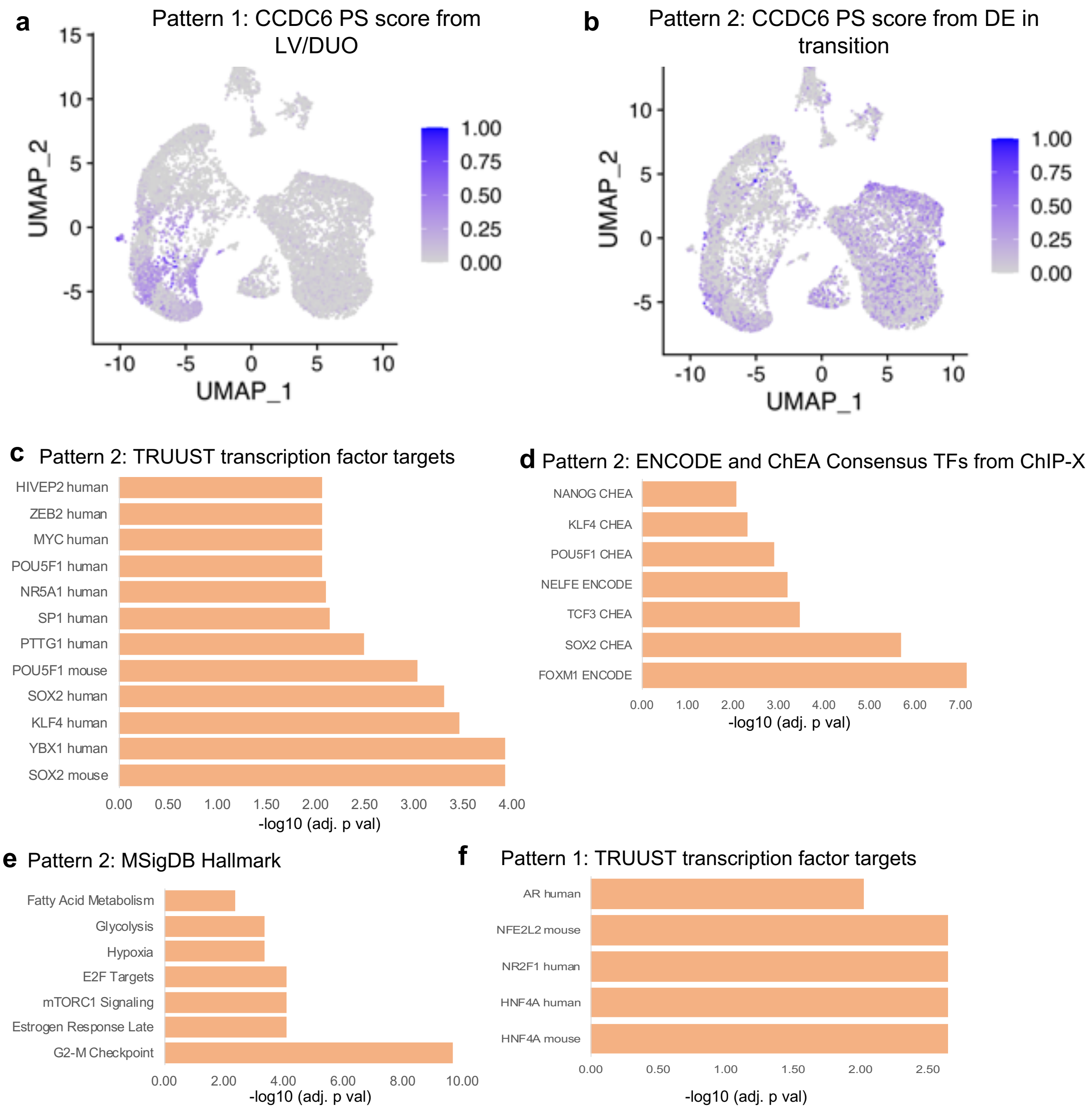


**Figure S10:** 10-clone scRNA-seq

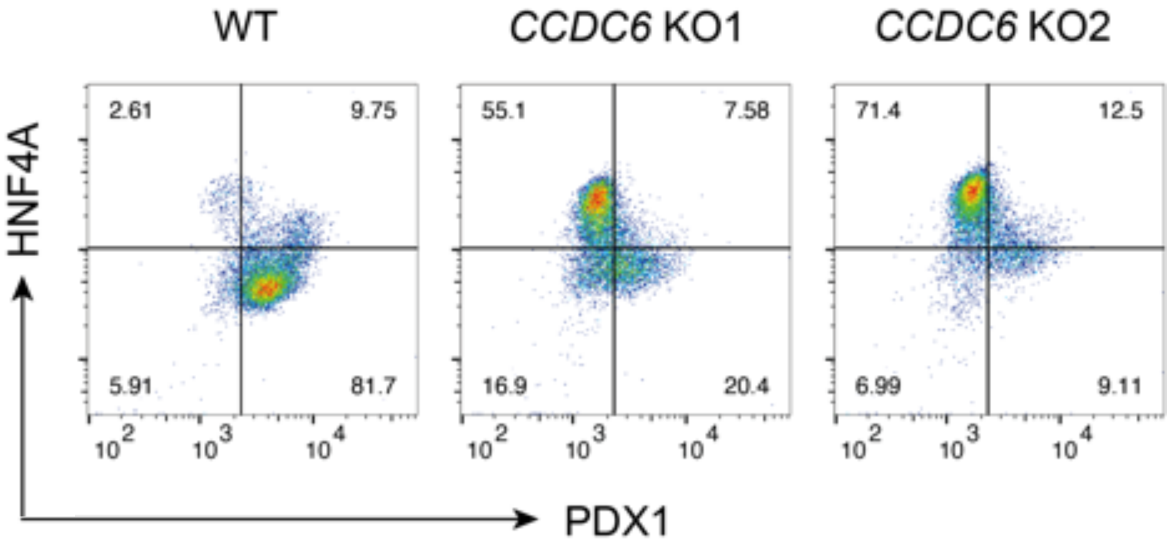


**Figure S11:** CCDC6 KO sorting.
